# Analysis of next- and third-generation RNA-Seq data reveals the structures of alternative transcription units in bacterial genomes

**DOI:** 10.1101/2021.01.02.425006

**Authors:** Qi Wang, Zhaoqian Liu, Bo Yan, Wen-Chi Chou, Laurence Ettwiller, Qin Ma, Bingqiang Liu

## Abstract

Alternative transcription units (ATUs) are dynamically encoded under different conditions or environmental stimuli in bacterial genomes, and genome-scale identification of ATUs is essential for studying the emergence of human diseases caused by bacterial organisms. However, it is unrealistic to identify all ATUs using experimental techniques, due to the complexity and dynamic nature of ATUs. Here we present the first-of-its-kind computational framework, named SeqATU, for genome-scale ATU prediction based on next-generation RNA-Seq data. The framework utilizes a convex quadratic programming model to seek an optimum expression combination of all of the to-be-identified ATUs. The predicted ATUs in *E. coli* reached a precision of 0.77/0.74 and a recall of 0.75/0.76 in the two RNA-Sequencing datasets compared with the benchmarked ATUs from third-generation RNA-Seq data. We believe that the ATUs identified by SeqATU can provide fundamental knowledge to guide the reconstruction of transcriptional regulatory networks in bacterial genomes.

## INTRODUCTION

An operon in bacterial genomes is defined as a group of consecutive genes regulated by a common promoter that all share the same terminator (*1*). Genes in the same operon generally encode proteins with relevant or similar biological functions; e.g., *lacZ, lacY*, and *lacA* in the *lac* operon encode proteins that help cells use lactose (*1, 2*). With decades of research on bacterial transcriptional regulation, the operon model has been found to have complex mechanisms that control expression (*3–5*). Multiple studies have shown that bacterial genes are dynamically transcribed under different triggering conditions, leading to shared genes among different mRNA transcripts (*6–8*). This dynamic architecture can be redefined by all of the alternative transcription units (a.k.a., ATUs) (*3, 5*), and more details can be found in fig. S1.

ATU identification is of fundamental importance for understanding the transcriptional regulatory mechanisms of bacteria, and these dynamic structures have been demonstrated to be associated with human diseases (*9–12*). For example, Bhat *et al*. studied the *alr-groEL1* operon, which is essential for the survival or virulence of *M. tuberculosis* (*9, 11*), the causative agent of tuberculosis (TB), and found that the regulation of the sub-operon is distinct from the main operon (*alr-groEL1* operon) under stress, especially during heat shock, pH, and SDS stresses (*9*). Another example is *Helicobacter pylori*, a gastric pathogen that is the primary known risk factor for gastric cancer (*12*). Sharma *et al*. found an acid-induced sub-operon *cag*22-18 transcribed from the primary *cag*25-18 operon in the *cag* pathogenicity island of the *H. pylori* genome under acid stress (*10*). The mechanism of the complex ATU structure in these pathogenic bacteria can help us to study the emergence of human diseases caused by bacterial organisms.

Several newly developed techniques have provided a comprehensive view of the *E. coli* transcriptome by identifying full-length primary transcripts (*13–17*). For example, SMRT-Cappable-seq (*6*) combines the isolation of the full-length bacterial primary transcriptome with PacBio SMRT (Single Molecule, Real-Time) sequencing (*6*), and simultaneous 5’ and 3’ end sequencing (SEnd-seq) (*7*) captures both transcription start sites (TSSs) and transcription termination sites (TTSs) via circularization of transcripts (*17*). Despite the great progress in experimental techniques, there are still some deficiencies. On the one hand, the read depth and error rate of the third-generation sequencing used in SMRT-Cappable-seq have an impact on ATU prediction compared with Illumina-based RNA-Seq (*7, 18*). On the other hand, the time-consuming, laborious, and costly properties of these experimental techniques make them unrealistic to be generally applicable to ATU predictions in bacteria under specific conditions. Thus, novel and robust computational methods for ATU identification in bacterial genomes based on RNA-Seq are urgently needed.

Fortunately, many computational studies have been carried out to predict ATUs in bacteria, which have provided some preliminary studies for ATU prediction. Several public databases, such as RegulonDB (*19*), DBTBS(*20*), MicrobesOnline (*21*), DOOR (*22, 23*), OperomeDB (*24*), DMINDA 2.0 (*25*), and ProOpDB (*26*), provide various levels of operon information and small amounts of ATU information. However, these databases cannot provide genome-scale ATU information under specific conditions. Some computational studies, including Rockhopper (*27*), SeqTU (*4, 28*), BAC-BROWSER(*29*), rSeqTU (*5*), and Operon-mapper (*30*), utilize machine learning and model integration methods based on genomic information and gene expression profiles to identify bacterial transcription architecture. However, these works still cannot solve the dynamic patterns and overlapping nature of ATUs.

Here, we present SeqATU, a novel computational method for genome-scale ATU prediction by analyzing next- and third-generation RNA-Seq data (Fig. 1 and table S1). SeqATU utilizes a convex quadratic programming model (CQP) and aims to provide the optimum expression combination of all of the to-be-identified ATUs. Specifically, CQP minimizes the squared error between the predicted expression level of ATUs and the actual expression levels in genetic and intergenic regions. It is noteworthy that SeqATU also utilizes the information about the bias rate function in modeling non-uniform read distribution as the linear constraints of CQP to profile the complexity of the ATU architecture. Overall, SeqATU provides a generalized framework for the inference of ATUs based on next-generation RNA-Seq data collected under multiple conditions and can be easily applied to any bacterial organism to identify the ATU architecture and construct a transcriptional regulatory network.

**Fig. 1.**
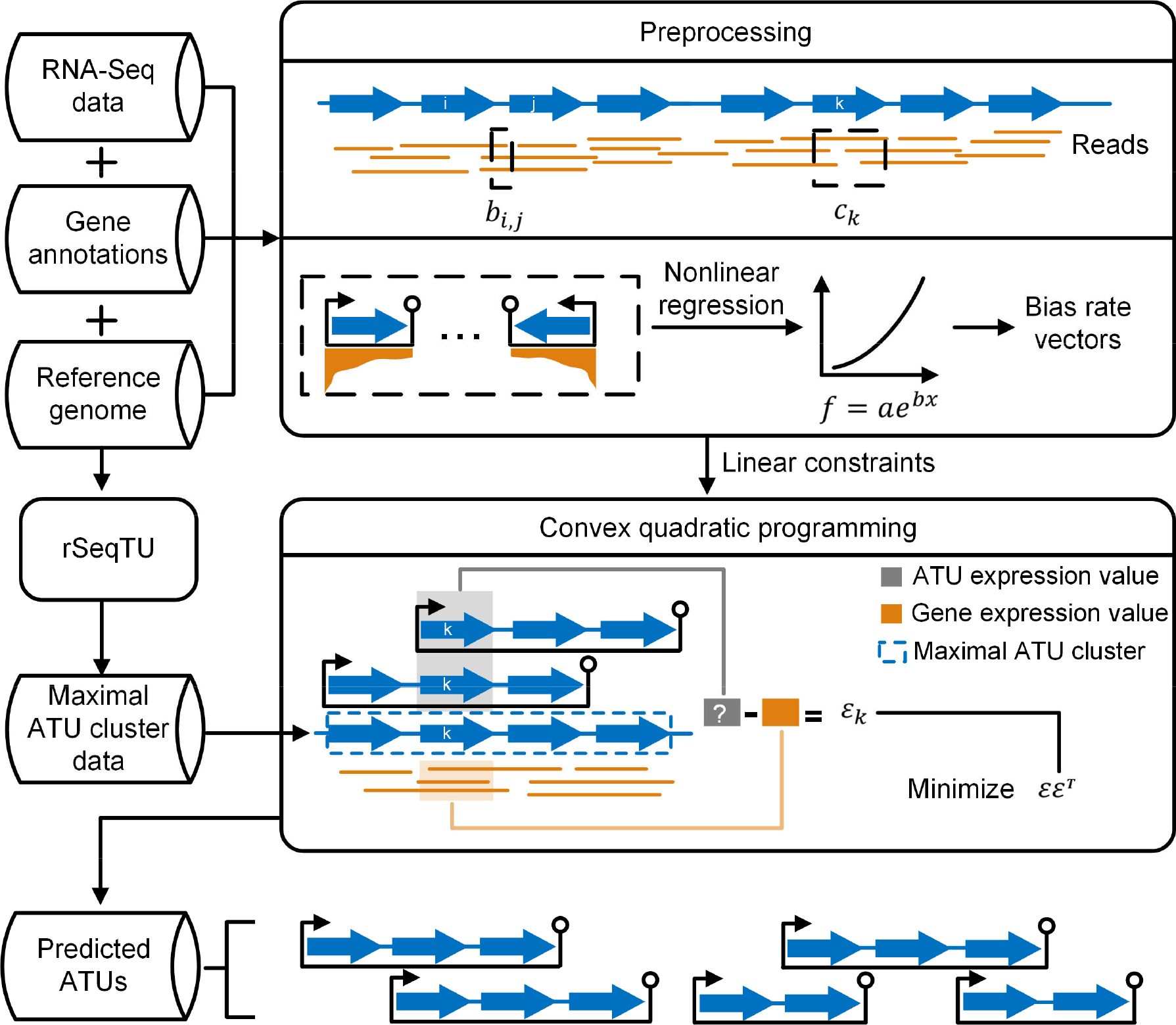
Schematic overview of SeqATU. The blue arrow and orange line denote gene and RNA-Seq read, respectively. The preprocessing stage requires RNA-Seq data in the FASTQ format, the reference genome sequence in the FASTA format, and gene annotations in the GFF format, generating linear constraints for the next convex quadratic programming (CQP) stage. There are two steps in the preprocessing stage: (*i*) calculating the expression value of the genetic region *c_k_* and intergenic region *b_i,j_* and (*ii*) modelling non-uniform read distribution along mRNA transcripts; specifically, we acquired a bias rate function *f*(*x*) = *ae^x^* using nonlinear regression and then constructed genetic or intergenic region bias rate vectors. The maximal ATU cluster data determined by rSeqTU and the linear constraints from preprocessing are both taken as inputs of CQP. CQP seeks the optimum expression combination of all of the to-be-identified ATUs to minimize the gap *εε^τ^* between the predicted ATU expression profile and the genetic and intergenic region expression profile. Finally, the output of CQP is the predicted ATUs.

## MATERIALS AND METHODS

### Data collection

The two Cappable RNA-Seq datasets used in this study, M9Enrich_Seq and RiEnrich_Seq, were obtained from *E. coli* grown under two different conditions: M9 minimal medium and Rich medium, respectively (6). The full-length primary transcripts were enriched as described in (6) with modifications to be adapted to Illumina sequencing. The capping and polyA tailing were performed as described in (6). The capped RNA was enriched using hydrophilic streptavidin magnetic beads (New England Biolabs) and eluted with Biotin using the same condition (6). Differently, the eluted RNA was enriched once more using streptavidin beads to further remove processed RNA (e.g., rRNA). Subsequently, the eluted RNA was used for library preparation using NEBNext Ultra II directional RNA library prep kit (E7760). Sequencing was performed on the Illumina Miseq system (paired-end, 100bp). All reads were mapped to the *E. coli* genome using Burrows-Wheeler Aligner (BWA) with the default parameters (*31*). Read alignment and other computational analyses were carried out using the *E. coli* genome NC_000913.3, and the corresponding gene annotations (GCF_000005845.2_ASM584v2_genomic.gff) were downloaded from NCBI. Two experimentally verified ATU datasets, SMRT_M9Enrich and SMRT_RiEnrich, were used as the benchmark data to evaluate the predicted ATUs, which were generated by SMRT-Cappable-seq under the same conditions as the Illumina datasets M9Enrich_Seq and RiEnrich_Seq, respectively (*6*). In addition, the ATUs defined by RegulonDB (*19*) and SEnd-seq (*7*) were also used as additional evaluation data in our study.

### Calculation of the expression values of genetic and intergenic regions

After the RNA-Seq reads in M9Enrich_Seq and RiEnrich_Seq were mapped to the *E. coli* genome using BWA, we determined the number of reads *N*(*l*) covering each genomic position *l*. Suppose that *g_i_* and *g*_*i*+1_ are two consecutive genes on the same strand; we denote the expression value of *g_i_* as *c_i_* and the expression value of the intergenic region between genes *g_i_* and *g*_*i*+1_ as *b*_*i,i*+1_. Then, the calculation of *c_i_* and *b*_*i,i*+1_ is given by:

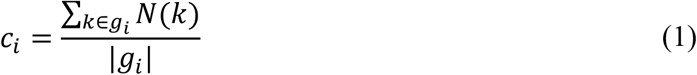

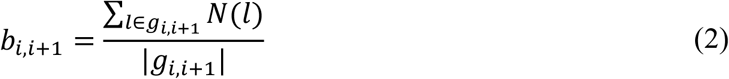

where *k* ∈ *g_i_* denotes that genomic position *k* is on the gene *g_i_* and |*g_i_*| denotes the genomic length of *g_i_*.

### Modeling non-uniform read distribution along mRNA transcripts

We introduced the bias rate function, which is similar to the bias curves in the work of Wu *et al*. (*32*), to address the non-uniform distribution of the RNA-Seq reads along mRNA transcripts (*32–35*). The bias function reflects the relative read distribution bias from the 3’ end to the 5’ end of an mRNA transcript. We assumed that the maximum read coverage of all the genomic positions of an mRNA transcript is the expression level without bias. It is noteworthy that a single gene mRNA transcript with no shared gene among different mRNA transcripts can serve as the ideal template for modeling non-uniform read distribution along mRNA transcripts. The specific steps of modeling non-uniform read distribution are detailed as follows:

#### Step 1: Single Gene mRNA Transcript Selection

We selected single gene mRNA transcripts from the evaluation data and plotted their expression distributions. Specifically, 12 groups of single gene mRNA transcripts with lengths ranging from 300 to 1,500 bp were selected from the evaluation data (more details are given in method S1), and each group had ten randomly chosen mRNA transcripts. Apparent decline trends appeared in the single gene mRNA transcripts with long lengths, ranging from 1,100 to 1,500 bp (fig. S2). The reason for this phenomenon may be that the incomplete transcription and 3’ end degradation or processing induce the enrichment of signal at 5’ end of the mRNA transcripts with long lengths (*36, 37*). Finally, we plotted the expression distribution of single gene mRNA transcripts with lengths ranging from 1,100 to 1,500 bp.

#### Step 2: Acquiring the Bias Rate Function

We applied nonlinear regression to the expression distribution of the selected single gene mRNA transcripts and acquired the hypothetical function *f*(*x*). Specifically, the *x* axis and *y* axis of the expression distribution were converted to the distance from the 3’ end of an mRNA transcript and the bias rate of read distribution, respectively. To apply nonlinear regression to single gene mRNA transcripts with different lengths, normalization was also implemented on *x*. Here, *x* = (*x*_1_, *x*_2_, …, *x_m_*) and *y* = (*y*_1_, *y*_2_, … *y_m_*) are defined by:

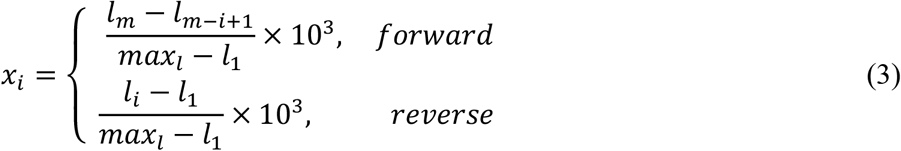

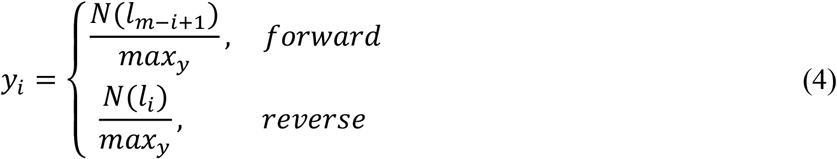

where *m* denotes the number of genomic positions on an mRNA transcript; *l* = (*l*_1_, *l*_2_, …, *l_m_*) denotes the genomic positions on an mRNA transcript; *max_l_* = *l_m_*; *N*(*l_i_*) denotes the expression level of the genomic position *l_i_*, i.e., the number of reads covering the genomic position *l_i_*; and *max_y_* denotes the expression level without bias in an mRNA transcript, which is calculated as *max* {*N*(*l_i_*)}, 1 ≤ *i* ≤ *m*. We used the function *nls* in R to acquire the hypothetical function *f*(*x*).

#### Step 3: Constructing Bias Rate Vectors

We constructed a genetic or intergenic region bias rate vector for each mRNA transcript by calculating the bias rate of all of its component genetic or intergenic regions. The bias rate of a genetic or an intergenic region is the average bias rate of all the genomic positions that it contains. Considering an mRNA transcript *T* and its component gene set {*g*_1_, *g*_2_, …, *g_n_*} (the details of the gene labels are described in method S2), we denoted the genetic region bias rate vector as ***u*** = (*u*_1_, *u*_2_, …, *u_n_*), which was calculated using the formula:

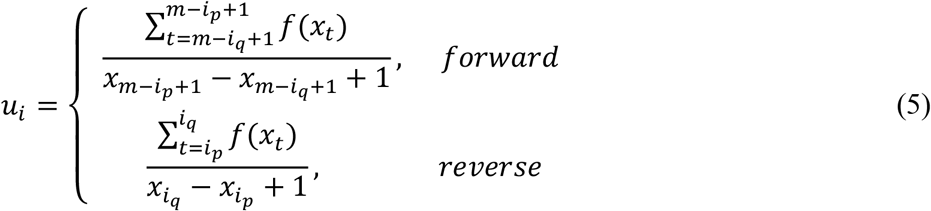

where *m* denotes the number of genomic positions on *T*; *u_i_* denotes the bias rate of *g_i_* for *T*; and *L_g_* = (*l*_1_*p*__, *l*_1_*q*__, *l*_2_*p*__,*l*_2_*q*__, …, *l_n_p__*, *l_n_q__*) is the range of the genomic positions of {*g*_1_, *g*_2_, …, *g_n_*}, while the range of the genomic positions of *g_i_* is [*l_i_p__, l_i_q__*], 1 ≤ *i* ≤ *n*. Similarly, the calculation of the intergenic region bias rate vector ***v*** = (*v*_1_, *v*_2_, …, *v*_*n*–1_) is provided in method S3.

### Modification of maximal ATU clusters

A maximal ATU cluster is defined as a maximal consecutive gene set such that each pair of its consecutive genes can be covered by at least one ATU. Similar to ATUs, maximal ATU clusters are also dynamically composed under different conditions or environmental stimuli in bacterial genomes (*5, 38*). Such a maximal ATU cluster can be used as an independent genomic region for ATU prediction, which alleviates the difficulty in computationally predicting ATUs at the genome scale. The output of our in-house tool rSeqTU can serve as the maximal ATU cluster data, which lays a solid foundation for ATU prediction (*5*). We modified the maximal ATU clusters from rSeqTU: (*i*) two maximal ATU clusters with distances less than 40 bp were combined into one cluster and (*ii*) a maximal ATU cluster was split at the intergenic region where the opposite-strand genes were located. In addition, we selected the maximal ATU clusters with expression values over ten (see the details in method S4), according to the study of Etwiller *et al*. (*13*).

### The mathematical programming model for ATU prediction

The predicted ATU expression profile should be consistent with the observed expression profiles of the genetic and intergenic regions. Therefore, the prediction of the ATU profiles can be modeled as an optimization problem, which seeks an optimum expression combination of all of the to-be-identified ATUs to minimize the gap between the predicted ATUs and the observed genetic and intergenic region expression profiles. Here, a convex quadratic programming model was built to solve this optimization problem.

We denoted a maximal ATU cluster as *G*, assuming that it contains the consecutive genes {*g*_1_, …, *g_n_*}, and the intergenic regions of these genes are {*g*_1,2_, …, *g*_*n*–1,*n*_}. The size of *G* is defined as the number of its component genes *n*. Theoretically, there are 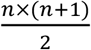 ATUs for *G*, and an ATU with consecutive genes {*g_i_*, *g*_*i*+1_, …, *g_j_*} is denoted as *a^i,j^*; the corresponding expression value is *x^i,j^*, 1 ≤ *i* ≤ *j* ≤ *n*.

For the component gene *g_k_* of *G*, the gap between the gene expression value *c_k_* and the sum of the expression level of the ATUs containing it is denoted as *τ_k_*, which provides the first *n* equality constraints in our mathematical programming model, *k* = 1,2, …, *n*. Similarly, for the intergenic region *g*_*l,l*+1_ of *G*, the gap between the intergenic region expression value *b*_*l,l*,+1_ and the sum of the expression level of the ATUs containing it is denoted as *β_l_*, providing the last *n* – 1 equality constraints in our mathematical programming model, *l* = 1,2, …, *n* – 1.

The goal of our mathematical programming model is to minimize the square of ***ε*** = (*τ*_1_, *τ*_2_, …, *τ_n_*, *β*_1_, …, *β*_*n*–1_), as the combination of *x^i,j^* with a minimal value of ***εε**^T^* is corresponding to an optimum expression combination of all ATUs *a^i,j^* for *G*, 1 ≤ *i* ≤ *j* ≤ *n*. Additionally, to control the number of optimal solutions and reduce the false-positive errors, we added an *L*^1^ regularization *α*||***x***||_1_ to ***εε**^T^* with *x^i,j^* ≥ 0, which is a linear function. Because of the variant expression level of different maximal ATU clusters, we used the expression value of *G* as *α*. In total, the convex quadratic programming model with unknown variables (***x, ε***) is shown as follows:

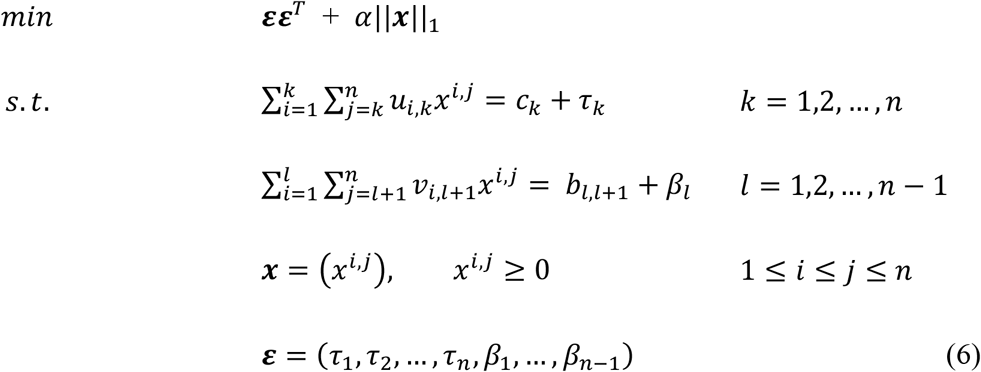

where ***u*** = (*u_i,j_*) is the genetic region bias rate vector for *G, u_i,j_* is the bias rate of gene *g_j_* for ATU *a^i,k^*, 1 ≤ *i* ≤ *j* ≤ *n*, *j* ≤ *k* ≤ *n*, ***v*** = (*v_p,q_*) is the intergenic region bias rate vector for *G*, and *v_p,q_* is the bias rate of the intergenic region *g*_*q*–1,*q*_ for ATU *a^p,l^*, 1 ≤ *p* < *q* ≤ *n*, *q* ≤ *l* ≤ *n* (see the details in method S5).

### Two evaluation methods for ATU prediction

In the first evaluation method, precision and recall were defined based on perfect matching (Eqs. 7). Perfect matching of two ATUs means that all of their component genes are the same. Here, the true positives (*TP*) are the number of predicted ATUs with the same component genes as an ATU in the evaluation data; the false positives (*FP*) are the number of predicted ATUs that do not exist in the evaluation data; the false negatives (*FN*) are the number of ATUs that appear in the evaluation data but not in the prediction results of SeqATU.

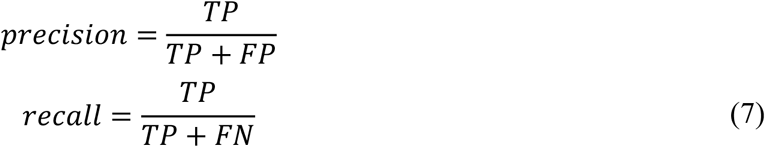

In the second evaluation method, precision and recall were defined based on relaxed matching, which is measured by the similarity of two ATUs. Assuming that an ATU *t* is in one of two datasets (the predicted ATU dataset and evaluated ATU dataset), the definition and calculation of the similarity of *t* are shown in the following three cases:

*Case 1*: If t shares boundary genes at both ends of an ATU in the other dataset, i.e., all component genes of *t* are the same as one in the other dataset, then *similarity*(*t*) = 1.

*Case 2*: If *t* shares exactly one boundary gene of ATUs in the other dataset, then we denote *U_a_* as the ATUs in the other dataset that share the 5’-end gene with *t* and denoted *U_b_* as the ATUs in the other dataset that share the 3’-end gene with *t*, 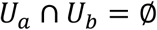, one of *U_a_* and *U_b_* can be empty. Then,

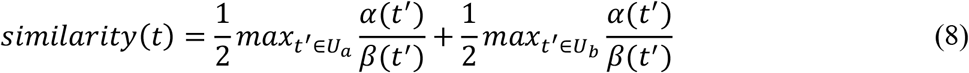

where *α*(*t′*) is the number of shared genes of *t* and *t*′ and *β*(*t′*) is the maximal size of *t* and *t*′.

*Case 3*: If *t* shares no boundary genes at both ends of the ATUs in the other dataset, then *similarity*(*t*) = 0.

Finally, the precision and recall based on relaxed matching are calculated by the following formula:

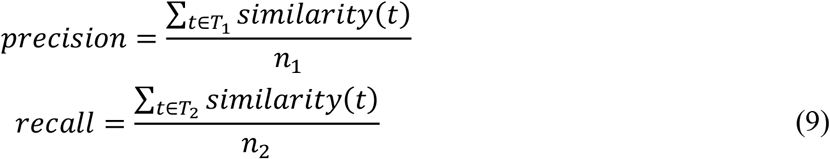

where *T*_1_ is the set of predicted ATUs, *n*_1_ is the number of predicted ATUs, *T*_2_ is the set of evaluated ATUs, and *n*_2_ is the number of evaluated ATUs.

## RESULTS

### A reliable bias rate function is acquired in modeling non-uniform read distribution along mRNA transcripts

To ensure the reliability of the bias rate function in modeling non-uniform read distribution, we selected four single gene mRNA transcript datasets randomly from the two evaluation datasets (SMRT_M9Enrich and SMRT_RiEnrich), named M9Enrich_1, M9Enrich_2, RiEnrich_1, and RiEnrich_2. Four bias rate functions, which are exponential functions, were generated after conducting nonlinear regression on the mRNA transcripts across these four datasets (Fig. 2). We found that these bias rate functions were similar (*R*^2^ > 0.998) when we evaluated the R-square statistic (for more details, see method S6 and table S2). The similarity of the four bias rate functions indicated that the selection of the single gene mRNA transcript datasets had little impact on modeling non-uniform read distribution along mRNA transcripts, implying the universal common non-uniform read distribution of different mRNA transcripts of *E. coli*. Specifically, we used the average of these four coefficients as the final coefficients of the exponential function, which was *f*(*x*) = *ae^bx^* with *a* = 0.256 and *b* = 0.00128.

**Fig. 2.**
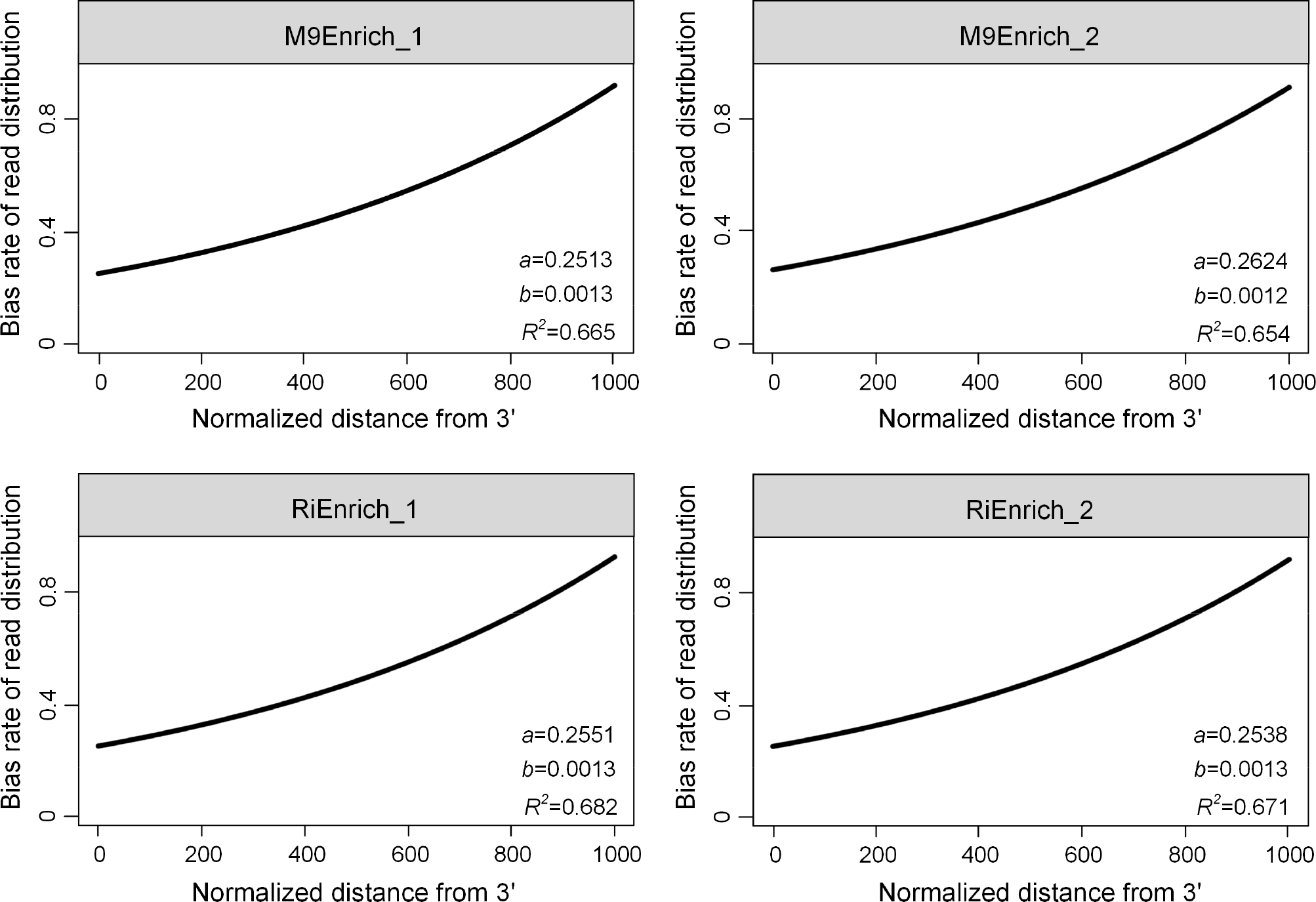
Results of modelling non-uniform read distribution along mRNA transcripts. The four bias rate functions (*y* = *ae^bx^*) by nonlinear regression had similar coefficients (*a* and *b*) across the four datasets M9Enrich_1, M9Enrich_2, RiEnrich_1 and RiEnrich_2.

### ATUs predicted by SeqATU reach precision and recall over 0.64

The performance evaluation was conducted by comparing the predicted ATUs with the ATUs in SMRT_M9Enrich and SMRT_RiEnrich, which were generated based on the third-generation sequencing and are not sensitive to transcripts with low expression levels. For a more accurate and fair evaluation, maximal ATU clusters after pre-selection were retained in the subsequent evaluations (more details about the pre-selection of maximal ATU clusters can be seen in method S7 and fig. S3).

The precision and recall of the predicted ATUs were calculated for each maximal ATU cluster. By considering only perfect matching, the average precision and recall were 0.67 and 0.67 for M9Enirch_Seq and 0.64 and 0.68 for RiEnrich_Seq, respectively. When using relaxed matching, the average precision and recall increased to 0.77 and 0.75 for M9Enrich_Seq and 0.74 and 0.76 for RiEnrich_Seq, respectively. The statistics for precision and recall on maximal ATU clusters with different sizes, as shown in Fig. 3A and fig. S4A. These results showed that the average precision and recall were decreasing with the increasing size of maximal ATU clusters (other than several large size ones due to their small number of counts). The results also indicated that the evaluation results based on relaxed matching were significantly higher than those based on perfect matching across different sizes. This result implied that the incorrectly predicted ATUs by SeqATU based on perfect matching tended to have strong similarities with the ATUs in the evaluation data. In addition, we also found that more than a quarter of the incorrectly predicted ATUs (25%/29% for M9Enrich_Seq/RiEnrich_Seq) by SeqATU based on perfect matching matched with the transcription units in RegulonDB (*19*).

**Fig. 3.**
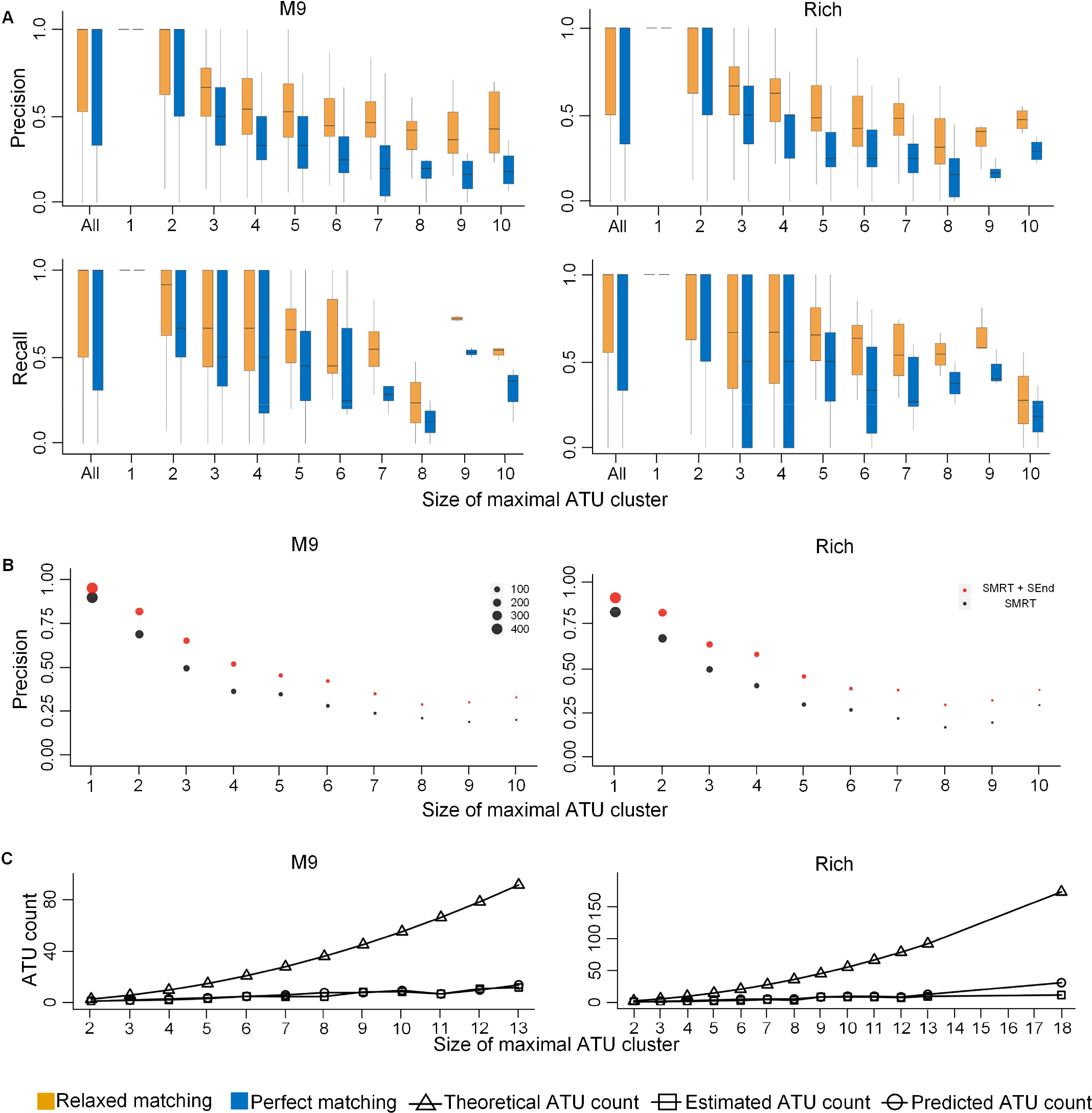
Overall evaluation results of SeqATU. (**A**) Precision and recall based on perfect matching and relaxed matching for M9Enrich_Seq (left) and RiEnrich_Seq (right) using evaluated ATUs from SMRT-Cappable-seq. (**B**) Average precision based on perfect matching for M9Enrich_Seq (left) and RiEnrich_Seq (right) using evaluated ATUs from SMRT-Cappable-seq (black) and evaluated ATUs from SMRT-Cappable-seq and SEnd-seq (red). The magnitude of the point denotes the number of maximal ATU clusters with same size. (**C**) Average number of ATUs across different sizes of SMRT maximal ATU clusters for M9Enrich_Seq (left) and RiEnrich_Seq (right).

The two evaluation datasets (SMRT_M9Enrich and SMRT_RiEnrich) were both from SMRT-Cappable-seq, while one of the processing steps of the technique filtered RNA reads smaller than 1,000 bp (*6*), which indicated that the ATUs in these two evaluation datasets were not comprehensive. To address this issue, we enriched the evaluation data by adding the ATUs defined by SEnd-seq (*7*), as SEnd-seq did not introduce any filtering based on RNA size. When we used the new evaluation data, the ATUs predicted by SeqATU improved by 15% (0.77) and 19% (0.76) in terms of the average precision based on perfect matching for M9Enrich_Seq and RiEnrich_Seq, respectively, and by 9% (0.84) and 12% (0.83) based on relaxed matching. The statistics for precision across different sizes of the maximal ATU clusters are shown in Fig. 3B and fig. S4B, showing that the values of precision based on perfect matching were significantly improved across different sizes of maximal ATU clusters by using the evaluated ATUs from SMRT-Cappable-seq and SEnd-seq. This result suggested that the ATUs we predicted, which were not in SMRT_M9Enrich and SMRT_RiEnrich, may be due to the RNA length selection of SMRT-Cappable-seq. We enriched the evaluation data by adding the ATUs in RegulonDB (*19*) and also found the improvement of precision across different sizes of maximal ATU clusters for M9Enrich_Seq and RiEnrich_Seq (fig. S4C).

Furthermore, to facilitate the understanding of the performance of SeqATU and to measure the influence of the maximal ATU clusters from rSeqTU on our ATU prediction method, SMRT maximal ATU clusters collected from SMRT_M9Enrich and SMRT_RiEnrich (for more details, see method S8) were applied for the CQP in two conditions (M9 minimal medium and Rich medium). We found that precision and recall increased to 0.73 and 0.77 for M9Enrich_Seq, respectively, and 0.69 and 0.80 for RiEnrich_Seq based on perfect matching (fig. S4D). Additionally, when using relaxed matching, precision and recall significantly increased to 0.82 and 0.84 for M9Enrich_Seq, respectively, and 0.79 and 0.86 for RiEnrich_Seq (fig. S4D). The significantly improved results verified the ability of SeqATU to accurately predict ATU when giving more accurate maximal ATU clusters. In addition, we found that the number of predicted ATUs and the evaluated ATUs under the maximal ATU cluster with the same size were similar except for the maximal size (Fig. 3C), and they were far less than the theoretical number, which indicated that SeqATU can effectively exclude most of the incorrect ATUs.

### The bias rate constraints efficiently improve the ability of SeqATU to predict ATUs

We tried to use SeqATU without bias rate constraints to predict the ATUs of *E. coli* and found that its performance significantly decreased compared with SeqATU (Fig. 4 and fig. S5). Specifically, the F-score of SeqATU without bias rate constraints was 0.69/0.68 based on perfect matching for M9Enrich_Seq/RiEnrich_Seq, compared with 0.75/0.74 for SeqATU. When using relaxed matching, the F-score of SeqATU without bias rate constraints was 0.79/0.78 for M9Enrich_Seq/RiEnrich_Seq, compared with 0.83/0.83 for SeqATU. This result suggested that the bias rate constraints of SeqATU could capture useful information about the non-uniform distribution of the RNA-Seq reads along the mRNA transcripts (*32–35*) and then efficiently improve the ability of the model to predict complex ATUs.

**Fig. 4.**
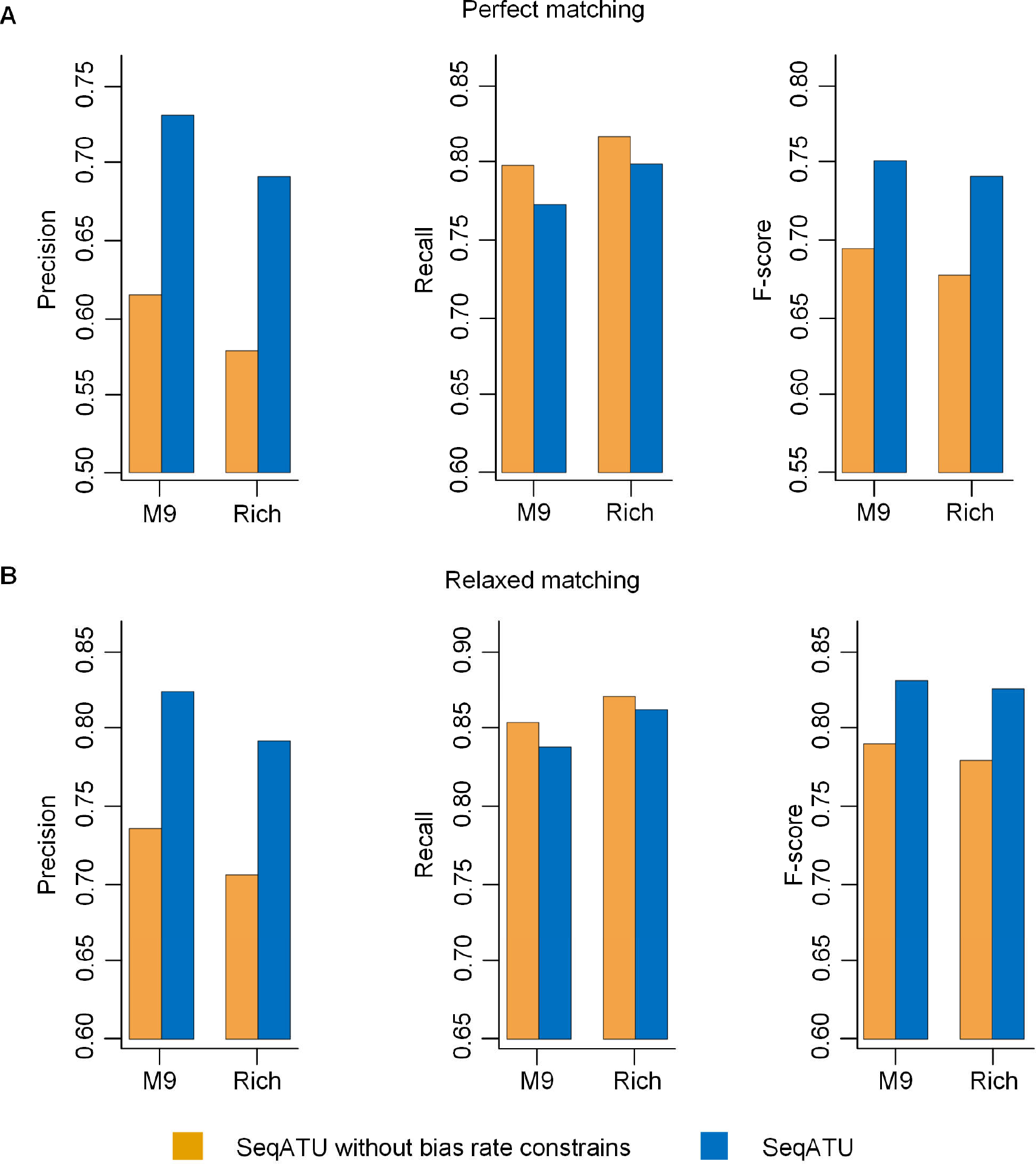
Comparative analysis of the performance between SeqATU and SeqATU without the bias rate constrains for SMRT maximal ATU clusters. (**A**) Precision, recall and F-score based on perfect matching for M9Enrich_Seq and RiEnrich_Seq. (**B**) Precision, recall and F-score based on relaxed matching for M9Enrich_Seq and RiEnrich_Seq.

### ATUs predicted by SeqATU display a dynamic composition and overlapping nature

A total of 2,973 distinct ATUs were identified in M9 minimal medium, and 2,767 were identified in Rich medium. Among them, there were 1,423/1,550 distinct ATUs on the forward strand and 1,323/1,444 on the reverse strand for M9Enrich_Seq/RiEnrich_Seq. Each of the predicted ATUs was comprised of an average of 2.59 genes, with the largest ATU containing 28 genes across the two conditions. The distribution of the size of the predicted ATUs is shown in Fig. 5A, from which we can see that the majority of ATUs (more than 87%) contained fewer than five genes in M9 minimal medium and Rich medium. Approximately 41% of the genes in *E. coli* were contained in more than one ATU for M9Enrich_Seq, compared to 43% genes for RiEnrich_Seq, suggesting that the ATUs in a maximal ATU cluster generally overlapped with each other (Fig. 5B). In addition, there were 1,576 ATU maximal clusters for M9Enrich_Seq and 1,512 ATU maximal clusters for RiEnrich_Seq. SeqATU identified a total of 1,977 identical ATUs under the two conditions, whereas there were 1,786 distinct ATUs. Among the distinct ATUs across the two conditions, 394 ATUs were from the same maximal ATU clusters in the two maximal ATU cluster datasets, and the rest were from different maximal ATU clusters. The fact there were distinct ATUs under the two conditions suggests that ATUs are dynamically responsive to different conditions or environmental stimuli (for more real examples about the ATUs under different conditions, see fig. S6).

**Fig. 5.**
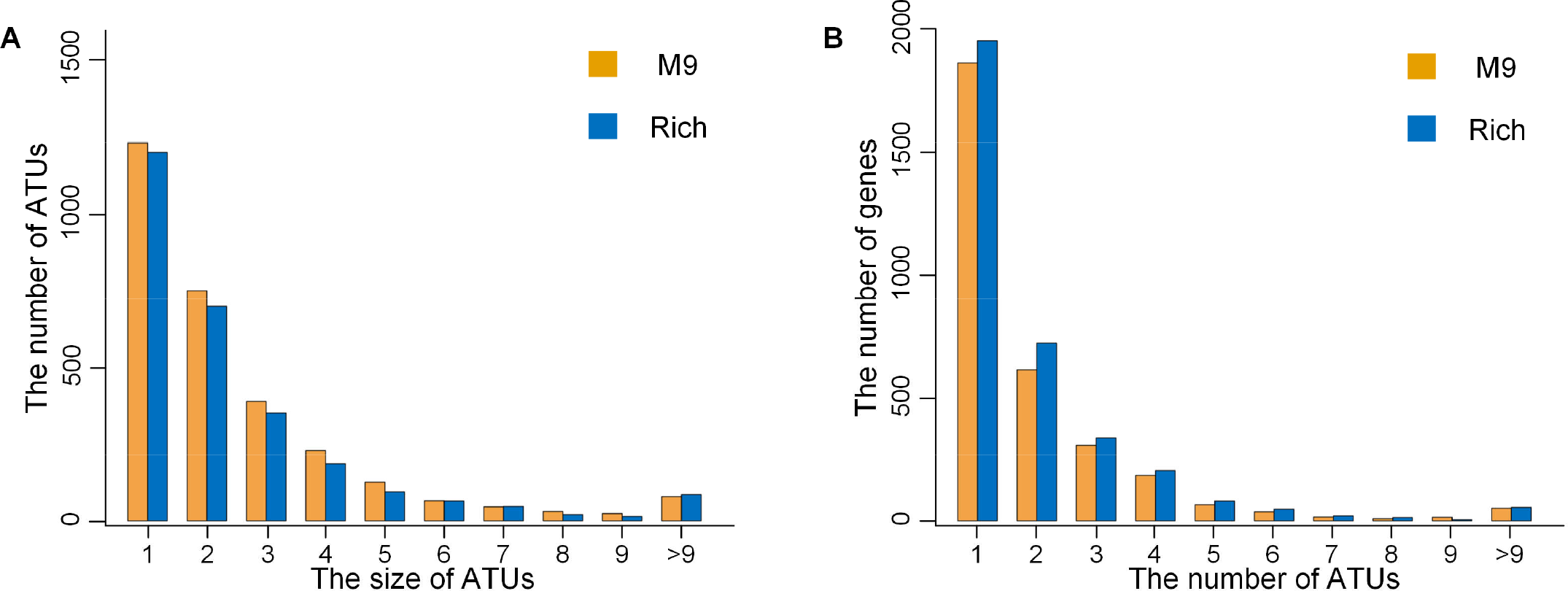
Comprehensive analysis of the predicted ATUs by SeqATU. (**A**) Number of ATUs across different sizes. The size of an ATU is the number of its component genes. (**B**) Distribution of the number of ATUs per gene.

The dynamic composition of predicted ATUs by SeqATU is of great significance to understand the interactions inside polymicrobial communities. For example, chronic airway infection by *Pseudomonas aeruginosa* considerably contributes to lung tissue destruction and impairment of pulmonary function in cystic-fibrosis (CF) patients (*39*). Marie *et al*. found that the presence of *E. coli* complemented the growth defect of a *P. aeruginosa bioA*-disrupted mutant that is unable to grow on rich medium, and can be beneficial to *P. aeruginosa* when biotin supply is limited (*39*). An ATU with a high expression level coded by the *uvrB* gene is identified by SeqATU in Rich medium, while it does not exist in M9 minimal medium (Fig. 6). We predicted the *uvrB* gene to be involved in the biotin metabolism pathway, as the *bioB, bioF, bioC*, and *bioD* genes contained in a same ATU with it have been known in the biotin metabolism KEGG pathway. Therefore, the observation by Marie *et al*. can be explained that the ATUs coded by the *uvrB* gene of *E. coli* can provide the biotin supply for *P. aeruginosa* under rich medium. This result showed that SeqATU could increase our understanding of interspecies competition and cooperation, which play an important role in shaping the composition and structure of polymicrobial bacterial populations.

**Fig. 6.**
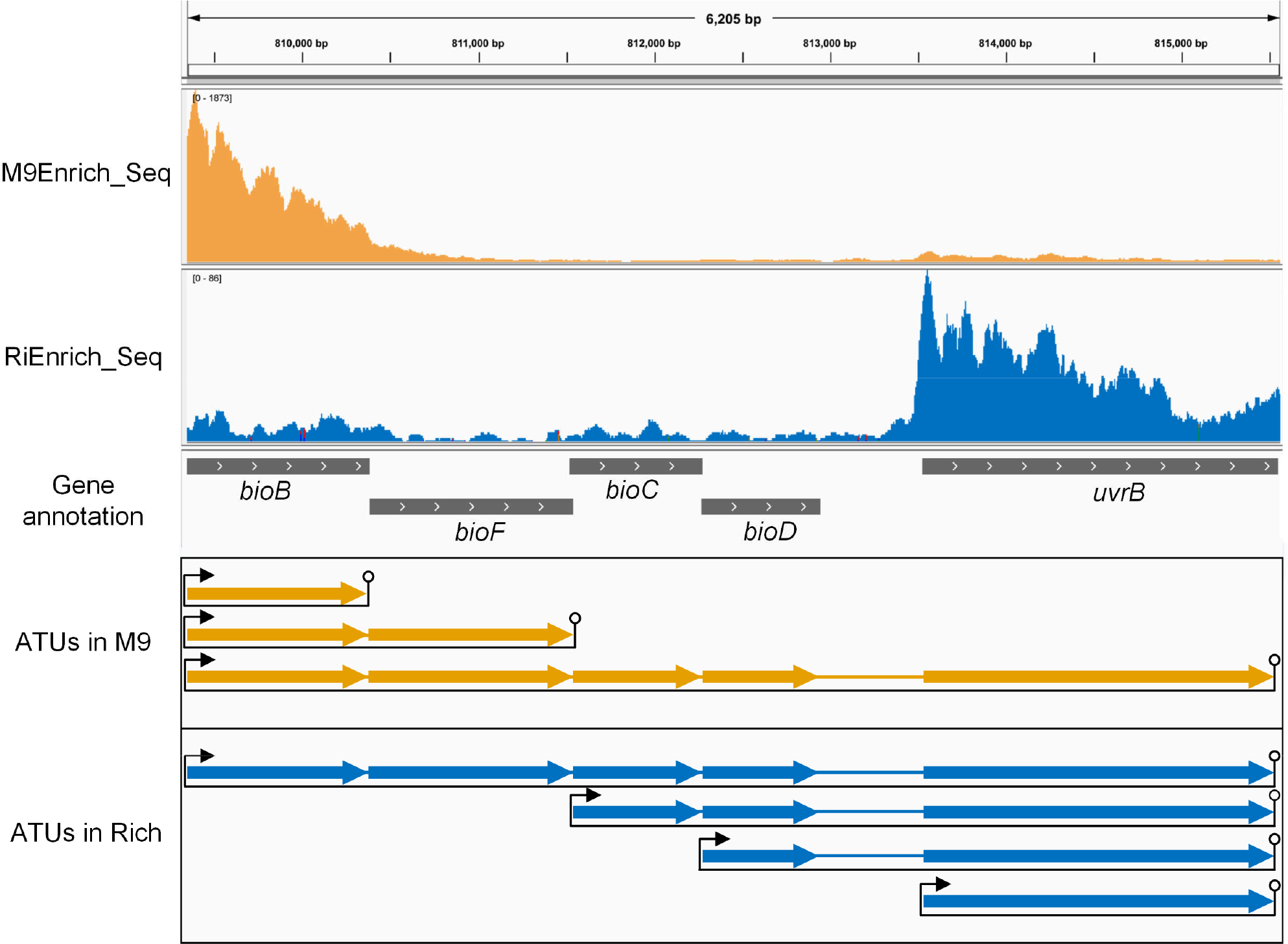
Integrative Genomics Viewer (IGV) representation of the mapping and ATUs. Mapping and ATUs of M9Enrich_Seq (orange) and RiEnrich_Seq (blue) were shown for the maximal ATU cluster containing the *bioB, bioF, bioC, bioD* and *uvrB* genes.

### Predicted ATUs by SeqATU are verified by experimental TSSs and TTSs

An experimental TSS dataset of *E. coli* from SEnd-seq (*7*) and a TF binding site dataset of *E. coli* from the experimental dataset of RegulonDB (*19*) were used to further verify the reliability of SeqATU and were named dataset 1 and dataset 2, respectively. There were 5,512 experimental TSSs in dataset 1 and 3,220 experimental TF binding sites in dataset 2. We considered the 5’-end genes and no 5’-end genes of the predicted ATUs by SeqATU. A gene that is not the 5’-end gene of any predicted ATU is named a no 5’-end gene. We identified 2,177/2,005 5’-end genes and 1,266/1,160 no 5’-end genes of the predicted ATUs for M9Enrich_Seq/RiEnich. A gene validated by experimental TSSs or TF binding sites means that it is the immediate downstream gene of an experimental TSS or TF binding site. As a result, the proportion of 5’-end genes of the predicted ATUs that were validated by experimental TSSs or TF binding sites was over 1.7 times greater than that of the no 5’-end genes (Table 1). Specifically, the proportion of 5’-end genes (29%/30% for M9Enrich_Seq/RiEnrich_Seq) validated by experimental TF binding sites was over three times greater than the no 5’-end genes (9.2%/9.0% for M9Enrich_Seq/RiEnrich_Seq). These results further verified the reliability of the ATUs predicted by SeqATU in terms of the TSS level. In addition, four other experimental TSS or promoter datasets from RegulonDB (*19*), dRNA-seq (*14*), and Cappable-seq (*13*) were also examined. The results are shown in table S3, and we also found a higher proportion of 5’-end genes of the predicted ATUs validated by experimental TSSs or promoters than that of no 5’-end genes.

**Table 1.** Results of predicted ATUs verified by experimental TSSs or TF binding sites. Overview of the experimental TSS and TF binding site datasets (dataset 1 and dataset 2) and the proportion of 5’-end genes and no 5’-end genes of the predicted ATUs by SeqATU for M9Enrich_Seq and RiEnrich_Seq, which were validated by experimental TSSs or TF binding sites.

|  |  | dataset 1 | dataset 2 |
| --- | --- | --- | --- |
| Source |  | Ju *et al.* (7) | RegulonDB TF binding sites |
| Technique |  | SEnd-seq | Collection |
| TSSs/TF binding sites |  | 5,512 | 3,220 |
| M9Enrich_Se<br>q | 5'-end genes | 83% | 29% |
|  | no 5'-end genes | 47% | 9.2% |
| RiEnrich_Seq | 5'-end genes | 89% | 30% |
|  | no 5'-end genes | 44% | 9.0% |

We also used two experimental TTS datasets of *E. coli* from SEnd-seq (*7*) and RegulonDB (*19*) to verify the reliability of predicted ATUs by SeqATU in terms of TTS level. These two experimental TTS datasets were named dataset 3 and dataset 4, respectively. There were 1,540 experimental TTSs in dataset 3 and 367 experimental TTSs in dataset 4. We considered the 3’-end genes and no 3’-end genes of the predicted ATUs by SeqATU. A gene that is not the 3’-end gene of any predicted ATU is named a no 3’-end gene. We identified 2,290/2,187 3’-end genes and 1,153/978 no 3’-end genes of the predicted ATUs for M9Enrich_Seq/RiEnrich_Seq. A gene validated by experimental TTSs means that it is the immediate upstream gene of an experimental TTS. As a result, the proportion of 3’-end genes of the predicted ATUs that were validated by experimental TTSs was over two times greater than that of no 3’-end genes (Table 2). Specifically, the proportion of 3’-end genes (51%/53% for M9Enrich_Seq/RiEnrich_Seq) validated by experimental TTSs from SEnd-seq was over three times greater than that of no 3’-end genes (15%/14% for M9Enrich_Seq/RiEnrich_Seq). These results further verified the reliability of the ATUs predicted by SeqATU in terms of the TTS level. In addition, two other computationally predicted TTS datasets from the works by Nadiras *et al*. (*40*) and Kingsford *et al*. (*41*) were also examined. The results are shown in table S4, and we also found the proportion of 3’-end genes (63%/62% for M9Enrich_Seq/RiEnrich_Seq) validated by computationally predicted Rho-independent TTSs was over two times greater than that of no 3’-end genes (29%/29% for M9Enrich_Seq/RiEnrich_Seq).

**Table 2.** Results of predicted ATUs verified by experimental TTSs. Overview of the experimental TTS datasets (dataset 3 and dataset 4) and the proportion of 3’-end genes and no 3’-end genes of the predicted ATUs by SeqATU for M9Enrich_Seq and RiEnrich_Seq, which were validated by experimental TTSs.

|  |  | dataset 3 | dataset 4 |
| --- | --- | --- | --- |
| Source |  | Ju *et al.* (7) | RegulonDB TTSs |
| Technique |  | SEnd-seq | Collection |
| TTSs |  | 1,540 | 3,67 |
| **M9Enrich_Se**<br>**q** | **3'-end genes** | 51% | 11% |
|  | **no 3'-end genes** | 15% | 5.2% |
| **RiEnrich_Seq** | **3'-end genes** | 53% | 11% |
|  | **no 3'-end genes** | 14% | 4.8% |

### The gene pairs frequently encoded in the same ATUs are more functionally related than those that can belong to two distinct ATUs

Functional analysis was conducted by integrating GO terms from the Gene Ontology (GO) database (*42*). In detail, we measured the level of functional relatedness for two types of consecutive gene pairs, which is similar to the definition in the work by Mao *et al*. (*38*). Two types of consecutive gene pairs were (*i*) gene pairs each consisting of a 5’-end gene of an ATU and the gene in its immediate upstream on the same strand and (*ii*) all the other gene pairs inside an ATU (Fig. 7A). In addition, we used a scoring scheme to measure the GO-based functional similarity between a pair of genes by Wu *et al*. (*43*). This study developed a GO similarity score and showed that the larger the score, the more likely that two genes are functionally related. In brief, the GO similarity score of a gene pair *g_k_* and *g_j_* is denoted as *S_GO_*(*g_k_, g_j_*):

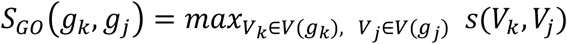

where *V_k_* and *V_j_* are the GO terms assigned to *g_k_* and *g_j_*, respectively; *s*(*V_k_*, *V_j_*) is the maximal number of common terms between paths in the two GO graphs induced by the GO terms *V_k_* and *V_j_*.

**Fig. 7.**
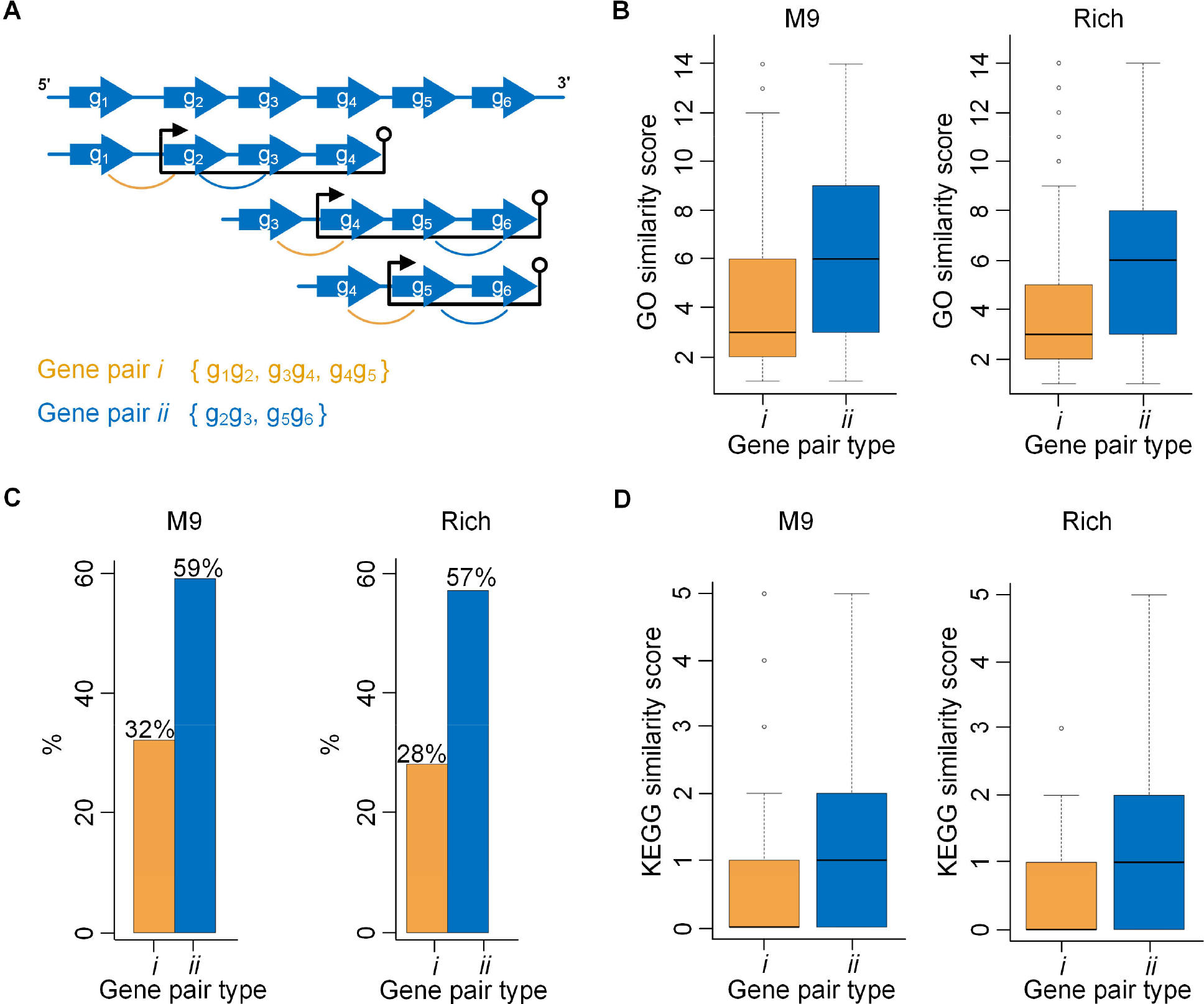
Interpretation and results of the functional relatedness of different gene pairs based on GO and KEGG enrichment analyses. (**A**) Illustration of two different gene pairs *i* and *ii*. (**B**) Functional relatedness results based on GO enrichment analysis for M9Enrich_Seq (left) and RiEnrich_Seq (right). (**C**) The proportion of two different gene pairs whose genes are contained in the same KEGG pathway for M9Enrich_Seq (left) and RiEnrich_Seq (right). (**D**) The functional relatedness results based on KEGG enrichment analysis for M9Enrich_Seq (left) and RiEnrich_Seq (right).

As a result, the mean GO similarity score was higher for type-*ii* gene pairs (5.97 *versus* 4.04 for M9Enrich_Seq and 5.86 *versus* 3.91 for RiEnrich_Seq) than for type-*i* gene pairs. A total of 574/524 type-*ii* gene pairs had GO similarity scores greater than four (64%/63% of a total of 899/834), while only 461/404 type-*i* gene pairs had GO similarity scores greater than four (36%/34% of a total of 1,274/1,179) for M9Enrich_Seq/RiEnrich_Seq. We also applied a *χ*^2^-test (*44*) to determine whether the distribution of *S_GO_*(*g_k_,g_j_*) was different for the type-*i* gene pairs and type-*ii* gene pairs. The *χ*^2^-statistics corresponded to a *P*-value less than 10^−4^, which revealed that the distribution of *S_GO_*(*g_k_, g_j_*) for the type-*ii* gene pairs was significantly different from the type-*i* gene pairs. Fig. 7B shows the distribution of *S_GO_* (*g_k_*, *g_j_*) for the type-*i* gene pairs and the type-*ii* gene pairs. These results strongly indicated that the type-*ii* gene pairs had a higher degree of GO similarity than the type-*i* gene pairs, suggesting that the gene pairs frequently encoded in the same ATUs (type-*ii* gene pairs) are more functionally related than those that can belong to two distinct ATUs (type-*i* gene pairs).

We also carried out a similar analysis of the two different gene pairs based on KEGG enrichment analysis (see more details in method S9) and found that the proportion of type-*ii* gene pairs (59%/57% for M9Enrich_Seq/RiEnrich_Seq), whose two genes were contained in the same KEGG pathway, was higher than the proportion of type-*i* gene pairs (32%/28% for M9Enrich_Seq/RiEnrich_Seq) (Fig. 7C). The distribution of the KEGG similarity scores of the two different types of gene pairs is shown in Fig. 7D, suggesting that genes of type-*ii* gene pairs have a higher probability of participating in the same KEGG pathway than those of type-*i* gene pairs.

## DISCUSSION

We developed SeqATU, the first computational method for genome-scale ATU prediction by analyzing next- and third-generation RNA-Seq data, using a CQP model. Linear constraints provided by the bias rate of read distribution were, for the first time, integrated into the CQP model. Positional bias refers to the non-uniform distribution of reads over different positions of a transcript (*33, 35*), which is handled by learning non-uniform read distributions from given RNA-Seq reads (*32*) or modeling the RNA degradation (*45*). The bias rate function we proposed can address the non-uniform read distribution along mRNA transcripts and also be desirable for standard next-generation RNA-Seq data that involves more degraded mRNAs, as the exponential function has been used to model the degradation of mRNA transcripts (*45*). As a result, a total of 2,973 distinct ATUs for M9Enrich_Seq and 2,767 distinct ATUs for RiEnrich_Seq were identified by SeqATU. The precision and recall reached 0.67/0.64 and 0.67/0.68, respectively, based on perfect matching and 0.77/0.74 and 0.75/0.76, respectively, based on relaxed matching for M9Enrich_Seq/RiEnrich_Seq. We further validated predicted ATUs using experimental transcription factor binding sites or transcription termination sites from RegulonDB and SEnd-Seq. In addition, the proportion of the 5’- or 3’-end genes of predicted ATUs that were validated by experimental transcription factor binding sites and transcription termination sites was over three times greater than that of no 5’- or 3’-end genes, demonstrating the high reliability of predicted ATUs. Gene pairs frequently encoded in the same ATUs were more functionally related than those that can belong to two distinct ATUs according to GO and KEGG enrichment analyses. These results demonstrated the reliability and accuracy of our predicted ATUs, implying the ability of SeqATU to reveal the transcriptional architecture of the bacterial genome.

In fact, the ATU architecture of bacteria is much more complex than that determined with currently used experimental techniques. We investigated the 5’-end genes and no 5’-end genes of the experimental ATUs identified by SMRT-Cappable-seq (*6*) using a combination of experimental TSSs from RegulonDB (*19*), dRNA-seq (*14*), Cappable-seq (*13*), and SEnd-seq (*7*). As a result, we found that the proportion of 5’-end genes (99%) validated by experimental TSSs was not significantly different from that of no 5’-end genes (92%). The high percentage of no 5’-end genes validated by experimental TSSs implied that the ATUs identified by experimental techniques are only a small proportion of the comprehensive ATUs in bacterial organisms due to the dynamic mechanisms of ATUs. These results further verified the necessity of developing robust computational methods for ATU identification.

SeqATU not only provides a powerful tool to understand the transcription mechanism of bacteria but also provides a fundamental tool to guide the reconstruction of a genome-scale transcriptional regulatory network. First, the ATU structure can help us to make new functional predictions, as genes in an ATU tend to have related functions. Second, ATUs can elucidate condition-specific uses of alternative sigma factors (*8, 46*). For example, the *thrLABC* operon is regulated by transcriptional attenuation. Totsuka *et al*. found that under the log phase growth condition, the *thrLABC* operon is the only transcript, while two transcripts are found under stationary phase growth condition, the *thrLABC* and *thrBC*. As validated experimentally, *σ^s^* can regulate the additional promoter located in front of *thrB* under the stationary phase growth condition and then separately regulate *thrBC*, which elucidates the condition-specific uses of *σ^s^* (*8*). Third, understanding the ATU structure is of great help to construct transcriptional and translation regulatory networks, such as for the construction of the σ-TUG (σ-factor-transcription unit gene) network (*47*). The transcription regulatory network consists of nodes (ATU and regulatory proteins) and links (interactions) (*48*), and the comprehensive ATU structure can provide a nearly complete set of nodes, which can improve the accuracy of regulatory prediction.

Although SeqATU has obtained satisfactory predicted results, there are still several challenges regarding the computational prediction of ATUs. On the one hand, due to the influence of the 3’ untranslated region (UTR) and 5’ untranslated region (UTR) in the intergenic regions, the expression value of intergenic regions cannot be reproduced perfectly by the same calculation used for the expression value of genetic regions. Without accurate reproduction, it is difficult to obtain the best expression combination of ATUs by the programming model based on the expression value of genetic and intergenic regions. On the other hand, due to the lack of strand-specific RNA-Seq data, it is difficult to distinguish the expression level of intergenic regions between two consecutive genes on the same strand derived from ATUs containing these two genes or antisense RNAs (asRNAs) (*6, 49*). All of these challenges and the great significance of ATU prediction inspire and encourage us to discover more information to determine the ATU structure in bacteria. For example, we plan to add high confidence TSSs and TTSs information to our programming model in the future. Additionally, since the microbiome is increasingly recognized as a critical component in human diseases, such as inflammatory bowel disease (*50*), antibiotic-associated diarrhoea (*51*), neurological disorders (*52*), and cancer (*53*) (*54*), predicting new ATUs of uncultured species from metagenomic and metatranscriptomic data is of great significance in uncovering new regulatory pathway and metabolic products during the development of diseases (*55*). However, due to a majority of species with unknown genomes or genome annotations within a microbial community, ATU prediction on metagenomics and metatranscriptomics is still a challenging task, which encourage us to pay more attention on it.

## Authors’ contributions

B.L., Q.M. and W.C. conceived the basic idea and designed the overall analyses. Q.W. carried out most of the computational analysis and data interpretation. All the authors wrote the manuscript.

## Data and materials availability

The raw data and source code of SeqATU and a detailed tutorial can be found at https://github.com/OSU-BMBL/SeqATU.

