## Supplementary material for "Analysis of next- and third-generation RNA-Seq data reveals the structures of alternative transcription units in bacterial genomes": Method S1-S9; Fig. S1-S6; Table S1-S4

**Method S1. Single gene mRNA transcripts selection**

We selected 12 groups of single gene mRNA transcripts with no shared gene from the evaluation data with lengths ranging from 300 to 1500 bp, each group has ten single gene mRNA transcripts selected randomly. The 12 groups of single gene mRNA transcripts are: the mRNA transcripts with lengths ranging from $l$ to $l+100$, $l = 300, 400, 500, 600, 700, 800, 900, 1000, 1100, 1200, 1300, 1400$.

**Method S2. Representation of gene labels**

Considering a maximal ATU cluster $G$ (or an mRNA transcript $G$), assuming that $G$ contains four consecutive component genes, and these four genes are labeled $g_1$, $g_2$, $g_3$, and $g_4$ from 5’ end to 3’ end of $G$. Besides, we denote $L_g = (l_{1_p}, l_{1_q}, l_{2_p}, l_{2_q}, l_{3_p}, l_{3_q}, l_{4_p}, l_{4_q})$ as the range of the genomic positions of $\{g_1, g_2, g_{3,}g_4\}$, concretely the range of the genomic positions of $g_i$ is $[l_{i_p}, l_{i_q}]$, $1 \le i \le 4$. And $L_{iter=}(l_{1_s}, l_{1_e}, l_{2_s}, l_{2_e}, l_{3_s}, l_{3_e})$ as the range of the genomic positions of $\{g_{1,2}, g_{2,3}, g_{3,4}\}$, concretely the range of the genomic positions of $g_{j,j+1}$ is $[l_{j_s}, l_{j_e}]$, $1 \le j \le 3$.

**Method S3. Calculation of the intergenic region bias rate vector**

For an mRNA transcript $T$ and its component intergenic region set $\{g_{1,2}, g_{2,3}, \dots, g_{n-1,n}\}$, we denote the intergenic region bias rate vector as $\boldsymbol{v} = (v_1, v_2, \dots, v_{n-1})$, which is calculated using the formula:

$$v_j = \begin{cases} \dfrac{\sum_{t=m-j_e+1}^{m-j_s+1} f(x_t)}{x_{m-j_s+1} - x_{m-j_e+1} + 1}, & forward \\ \dfrac{\sum_{t=j_s}^{j_e} f(x_t)}{x_{j_e} - x_{j_s} + 1}, & reverse \end{cases} \tag{1}$$

where $m$ denotes the number of genomic positions on $T$, $v_j$ denotes the bias rate of $g_{j,j+1}$ for $T$, $L_{iter=}(l_{1_s}, l_{1_e}, l_{2_s}, l_{2_e}, \dots, l_{n-1_s}, l_{n-1_e})$ is the range of the genomic positions of $\{g_{1,2}, g_{2,3}, \dots, g_{n-1,n}\}$, and the range of the genomic positions of $g_{j,j+1}$ is $[l_{j_s}, l_{j_e}]$, $1 \le j \le n-1$.

**Method S4. Calculation of the expression value of a maximal ATU cluster**

For a maximal ATU cluster $G$, the expression value of $G$ is:

$$\frac{\sum_{k \in G} N(k)}{|G|} \tag{2}$$

where $|G|$ is the genomic length of $G$, i.e. the number of genomic positions on $G$; $k \in G$ denotes that the genomic position $k$ is on $G$; $N(k)$ is the number of reads covering the position $k$ on the genome.

**Method S5. Combination of genetic/intergenic region bias rate vectors**

Assume that a maximal ATU cluster $G$ contains consecutive genes $\{g_1, g_2, g_3\}$ and the intergenic regions $\{g_{1,2}, g_{2,3}\}$, then there are six ($\frac{3\times(3+1)}{2} = 6$) ATUs with different consecutive genes for $G$, which are $a^{1,1}$, $a^{1,2}$, $a^{1,3}$, $a^{2,2}$, $a^{2,3}$, and $a^{3,3}$. Concretely, we merged the genetic region bias rate vectors of ATUs with the same 5'-end gene $g_1$, $a^{1,1}$, $a^{1,2}$, and $a^{1,3}$, into one genetic region bias rate vector $(u_{1,1}, u_{1,2}, u_{1,3})$, i.e., the genetic region bias rate vector of ATU $a^{1,3}$ which has the maximal size among $a^{1,1}$, $a^{1,2}$, and $a^{1,3}$. Likewise, the genetic region bias rate vectors of ATUs with the same 5'-end gene $g_2$, $a^{2,2}$ and $a^{2,3}$, were merged into the genetic region bias rate vector $(u_{2,3}, u_{33})$, i.e., the genetic region bias rate vector of $a^{2,3}$. And the genetic region bias rate vector with the same 5'-end gene $g_3$, $a^{3,3}$, was merged into $(u_{3,3})$. Finally, the genetic region bias rate vector for $G$ is $\boldsymbol{u} = (u_{1,1}, u_{1,2}, u_{1,3}, u_{2,2}, u_{2,3}, u_{3,3})$.

Similarly, we merged the intergenic region bias rate vectors of ATUs with the same 5'-end gene $g_1$, $a^{1,2}$ and $a^{1,3}$, into one bias rate vector $(v_{1,2},\ v_{1,3})$, i.e. the intergenic region bias rate vector of $a^{1,3}$; the intergenic region bias rate vectors of ATUs with the same 5'-end gene $g_2$, $a^{2,3}$, were merged into $(v_{2,3})$. The intergenic region bias rate vector for $G$ is $\boldsymbol{v} = (v_{1,2}, v_{1,3}, v_{2,3})$.

**Method S6. Similarity of four bias rate functions obtained by nonlinear regression**

To measure the similarity of the four bias rate functions $f_1(x) = 0.251319e^{0.001298x}$, $f_2(x) = 0.262360e^{0.001245x}$, $f_3(x) = 0.255069e^{0.001291x}$, and $f_4(x) = 0.253783e^{0.001286x}$, we evaluated the R-square statistic. Concretely, as for $f_j(x)$, $j = 1,2,3,4$, we calculated the $R^2$ between function $f_j(x)$ and the dataset containing the value of $f_i(x)$ with $x$ ranging from 1 to 1,000, $i \neq j$, $i = 1,2,3,4$. The results of the R-square statistic were showed in Supplementary Table 4.

**Method S7. Selection of maximal ATU clusters**

Maximal ATU clusters whose component genes were not involved in the ATUs identified by SMRT-

Cappable-seq were not considered in the performance evaluation section. 501/534 (32%/35% of a total of 1,576/1,512) maximal ATU clusters were not considered in evaluation section for M9Enrich_Seq/RiEnrich_Seq. 64%/58% of these maximal ATU clusters possess low expression value (<40), due to the insensitivity of third-generation sequencing for transcripts with low expression levels. While 36%/42% possess high expression value, which may be caused by the deficiency of SMRT-Cappable-seq for identifying ATUs. More examples of Integrative Genomics Viewer (IGV) representation for maximal ATU clusters with high expression value yet not identified by SMRT-Cappable-seq were showed in fig. S6.

**Method S8. Collection of the SMRT maximal ATU clusters from SMRT_M9Enrich and SMRT_RiEnrich**

We collected ATUs from SMRT_M9Enrich and SMRT_RiEnrich to acquire the SMRT maximal ATU clusters under M9 minimal medium and Rich medium, respectively. All ATUs that overlap with each other are in the same cluster, which also means ATUs that belong to different clusters cannot overlap with each other. At last, the consecutive genes in a same cluster make up an SMRT maximal ATU cluster.

**Method S9. KEGG enrichment analysis of type-*i* and type-*ii* gene pairs**

For the gene pairs, type-*i* and type-*ii* gene pairs, we considered the gene pairs whose genes are contained in the same KEGG pathway. We defined the KEGG similarity score of a gene pair as the number of same KEGG pathways which its two genes are contained.

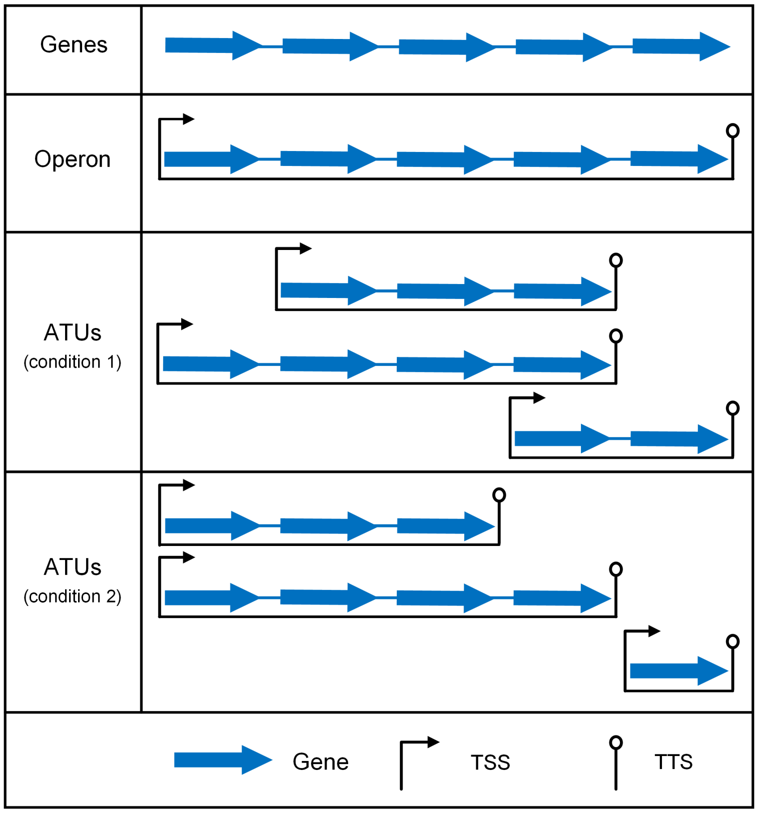


**Fig. S1. A diagram of operon and ATUs.** ATUs are dynamically composed of different consecutive genes under condition 1 and 2, and may overlap with each other.

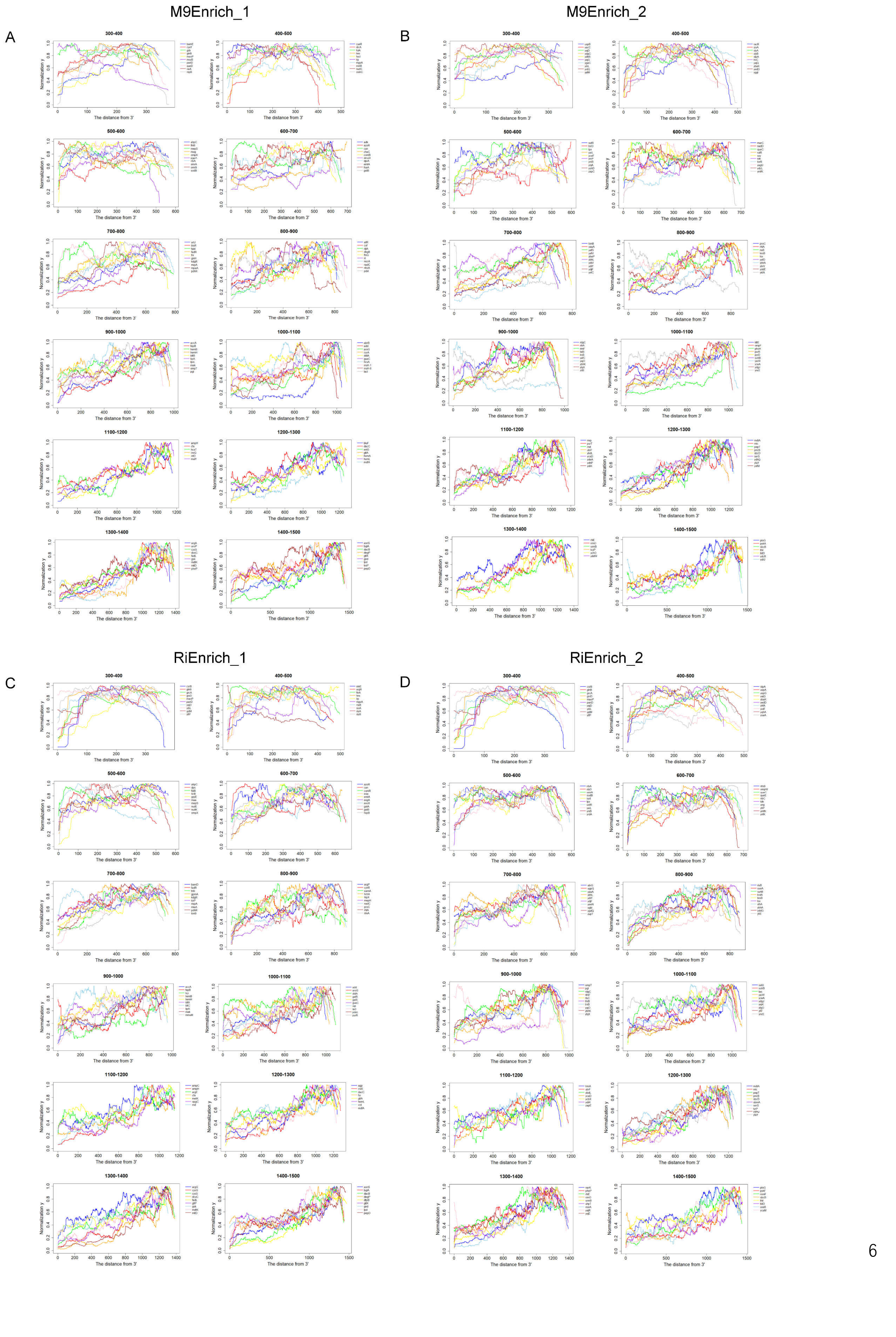

M9Enrich_1
A
M9Enrich_2
B
RiEnrich_1
C
RiEnrich_2
D
300-400
400-500
500-600
600-700
700-800
800-900
900-1000
1000-1100
1100-1200
1200-1300
1300-1400
1400-1500
Normalization y
The distance from 3'

**Fig. S2. The expression distribution of single gene mRNA transcripts.** (**A**) The expression distribution of the single gene mRNA transcripts across different length groups of M9Enrich_1. (**B**) The expression distribution of the single gene mRNA transcripts across different length groups of M9Enrich_2. (**C**) The expression distribution of the single gene mRNA transcripts across different length groups of RiEnrich_1. (**D**) The expression distribution of the single gene mRNA transcripts across different length groups of RiEnrich_2.

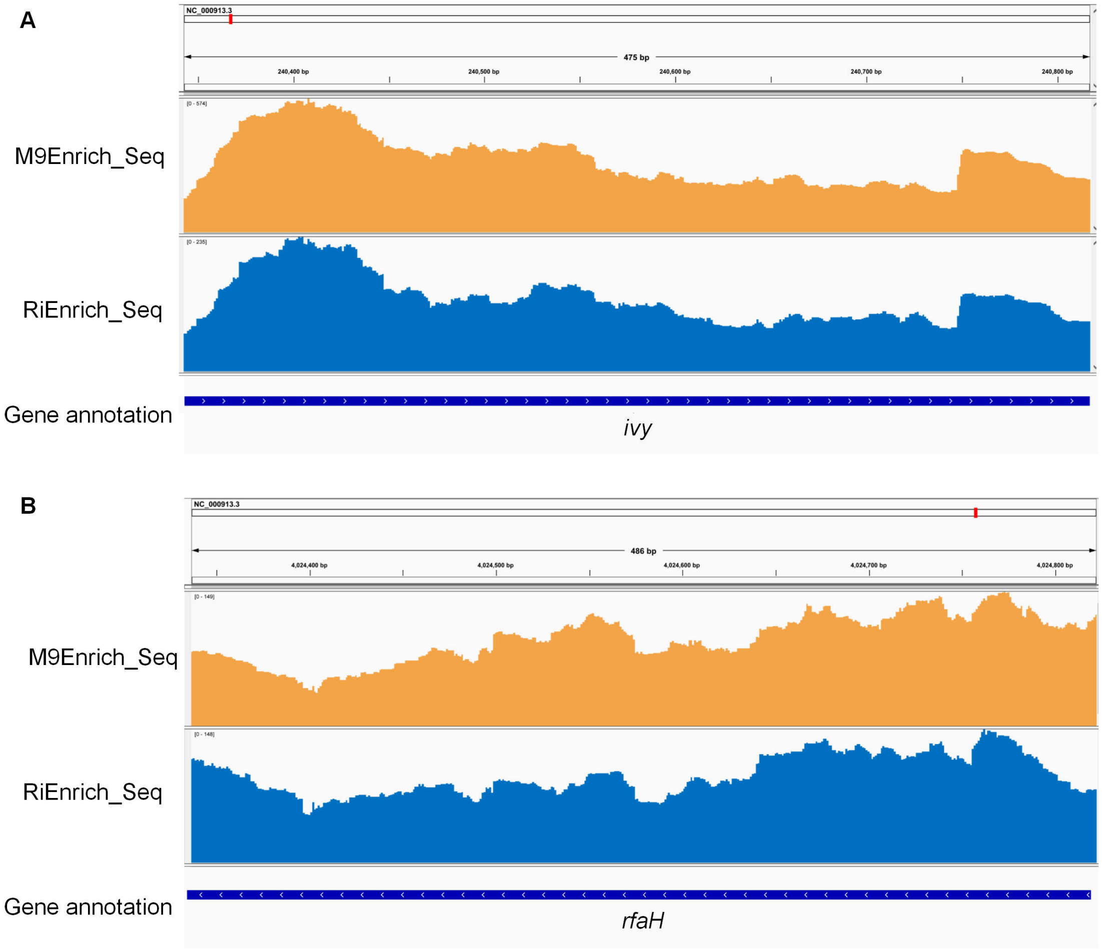


**Fig. S3. Integrative Genomics Viewer (IGV) representation of the mapping for maximal ATU**

**clusters.** IGV representation of two maximal ATU clusters with high expression value, which were from rseqTU yet are not identified by SMRT-Cappable-seq. (**A**) IGV representation of the mapping for the maximal ATU cluster containing the *ivy* gene with expression value 371/141 for M9Enrich_Seq/RiEnrich_Seq. (**B**) IGV representation of the mapping for the maximal ATU cluster containing the *rfaH* gene with expression value 125/120 for M9Enrich_Seq/RiEnrich_Seq.

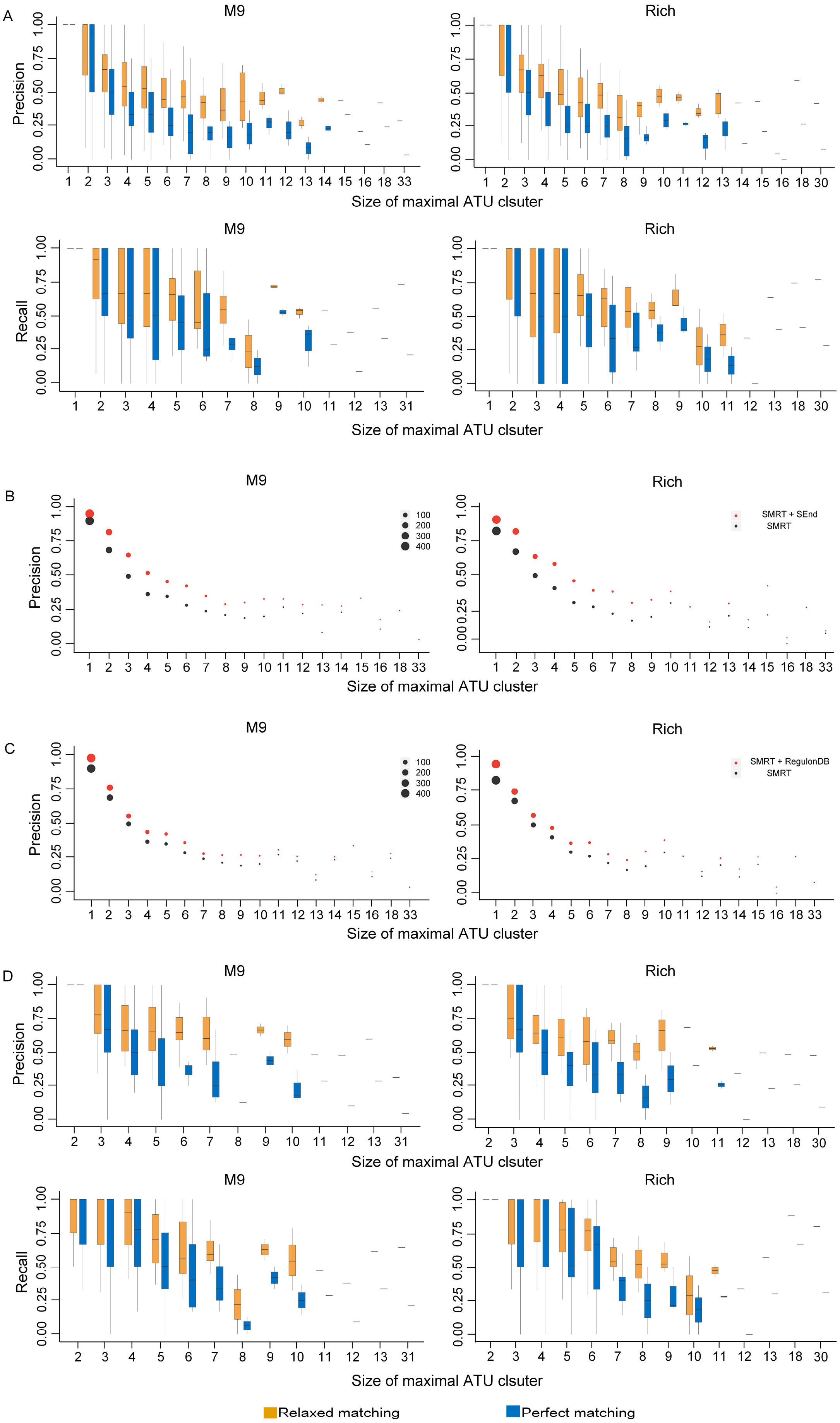
A
M9
Rich
Precision
Size of maximal ATU clsuter
Recall
B
M9
Rich
Precision
100
200
300
400
SMRT + SEnd
SMRT
Size of maximal ATU cluster
C
M9
Rich
Precision
100
200
300
400
SMRT + RegulonDB
SMRT
Size of maximal ATU cluster
D
M9
Rich
Precision
Size of maximal ATU clsuter
Recall
Size of maximal ATU clsuter
Relaxed matching
Perfect matching

**Fig. S4. Overall evaluation results of SeqATU.** (**A**) Precision and recall based on perfect matching and relaxed matching across different sizes of maximal ATU clusters for M9Enrich_Seq (left) and RiEnrich_Seq (right). (**B**) Average precision based on perfect matching across different sizes of maximal ATU clusters for M9Enrich_Seq (left) and RiEnrich_Seq (right) using evaluated ATUs from SMRT-Cappable-seq (black) and evaluated ATUs from SMRT-Cappable-seq and SEnd-seq (red). The magnitude of the point denotes the number of maximal ATU clusters with same size. (**C**) Average precision based on perfect matching across different sizes of maximal ATU clusters for M9Enrich_Seq (left) and RiEnrich_Seq (right) using evaluated ATUs from SMRT-Cappable-seq (black) and evaluated ATUs from SMRT-Cappable-seq and RegulonDB (red). (**D**) Precision and recall based on perfect matching and relaxed matching across different sizes of maximal ATU clusters for M9Enrich_Seq(left) and RiEnrich_Seq(right) using SMRT maximal ATU clusters.

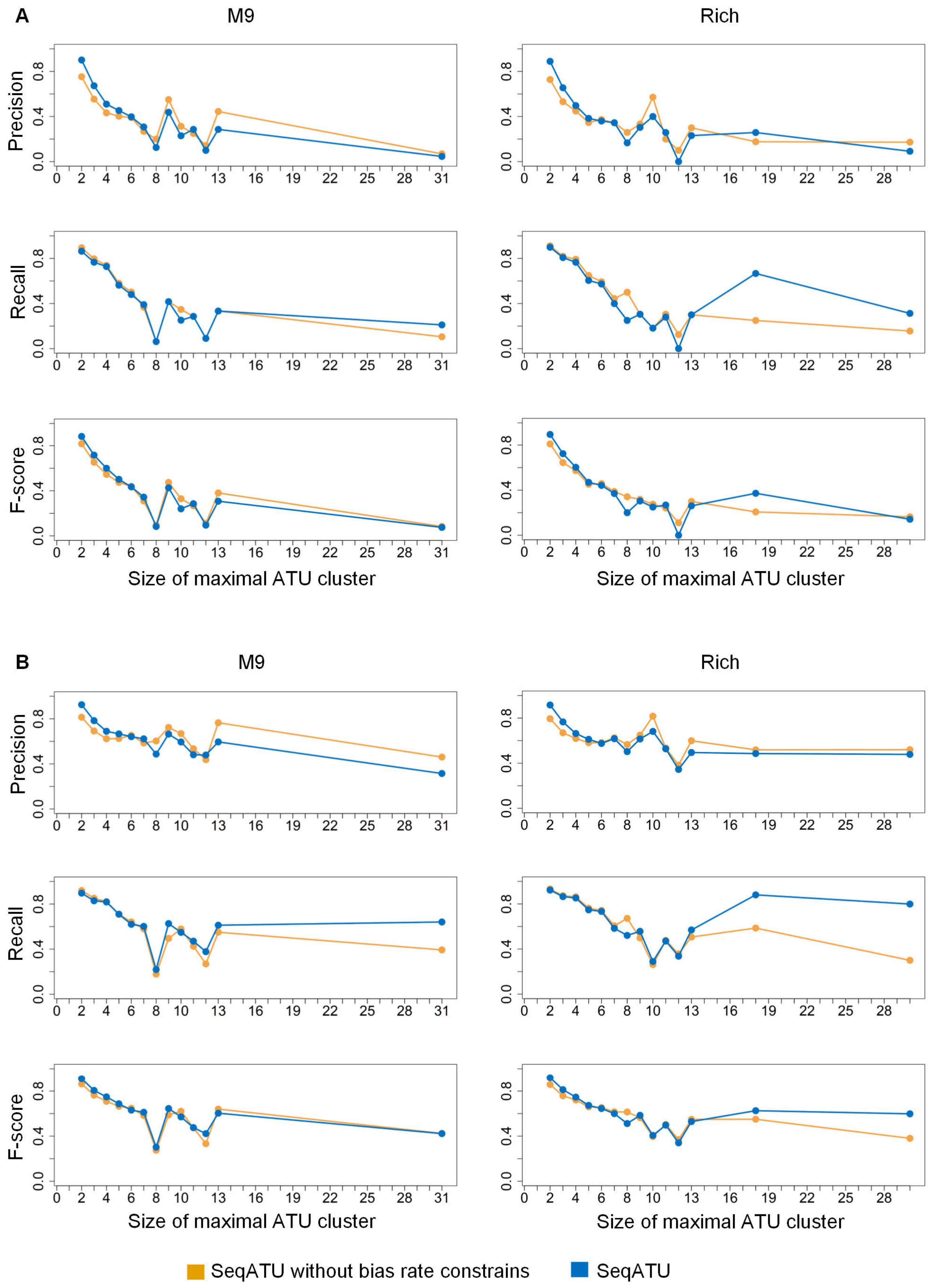
A
M9
Rich
Precision
Recall
F-score
Size of maximal ATU cluster
B
M9
Rich
Precision
Recall
F-score
Size of maximal ATU cluster
SeqATU without bias rate constrains
SeqATU

99

**Fig. S5. Comparative analysis of the performance between SeqATU and SeqATU without bias rate constrains across different sizes of maximal ATU clusters.** (**A**) Precision, recall, and F-score based on perfect matching for M9Enrich_Seq and RiEnrich_Seq. (**B**) Precision, recall, and F-score based on relaxed matching for M9Enrich_Seq and RiEnrich_Seq.

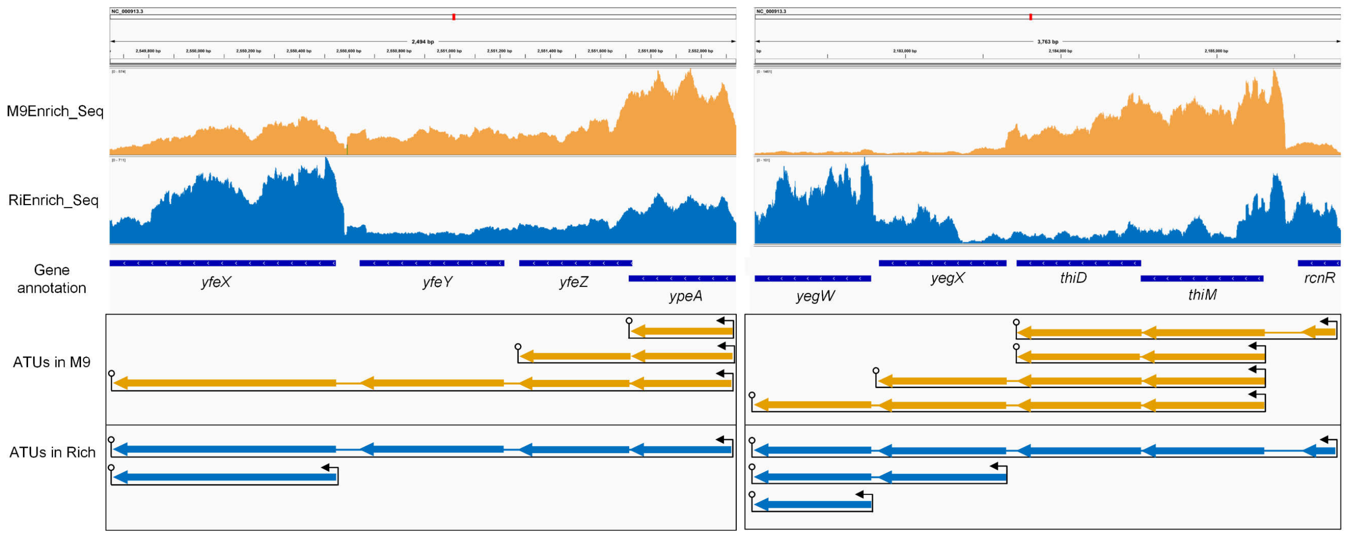


**Fig. S6. Integrative Genomics Viewer (IGV) representation of the mapping and ATUs.** IGV representation of ATUs in M9 minimal medium (orange) compared to Rich medium (blue) for the maximal ATU cluster containing the *yfeX*, *yfeY*, *yfeZ*, and *ypeA* genes (left) and the maximal ATU cluster containing the *yegW*, *yegX*, *thiD*, *thiM*, and *rcnR* genes.

**Table S1. Pseudo code of SeqATU.**

| **Algorithm:** SeqATU |
| --- |
| **Input:** Maximal ATU cluster data and RNA-Seq data |

**Output:** Predicted ATUs

**for** $G$ **in** maximal ATU cluster data **do**

**Preprocessing**:

$n \leftarrow$ the size of $G$

**Initialize** $L$ as the genomic positions of $G$

**Initialize** $L_g$ as the range of genomic positions of genetic regions of $G$

**Initialize** $L_{iter}$ as the range of genomic positions of intergenic regions of $G$

$B, C \leftarrow$ EXPRESSIONVALUE$(L_g, L_{iter}, n)$;

$u, v \leftarrow$ BIASRATEVECTOR$(G, L, L_g, L_{iter})$;

**Convex quadratic programming:**

**Input** $G$, $B$, $C$, $u$, and $v$ **to** convex quadratic programming;

**Output** the expression value $x$ of all ATUs for $G$

**end for**

**Function** EXPRESSIONVALUE$(L_g, L_{iter}, n)$ **begin**

**Initialize** $C$ and $B$ as empty arrays;

**for** $i \leftarrow 1$ **to** $n$ **do**

$sum = 0$;

**for** $j \leftarrow l_{i_p}$ **to** $l_{i_q}$ **do**

$sum = sum + N(j)$;

**end for**

$c_i = sum/(l_{i_q} - l_{i_p} + 1)$;

Append $c_i$ to $C$;

**end for**

**for** $i \leftarrow 1$ **to** $n - 1$ **do**

$sum = 0$;

**for** $j \leftarrow l_{i_s}$ **to** $l_{i_e}$ **do**

$sum = sum + N(j)$;

**end for**

$b_{i,i+1} = sum/(l_{i_e} - l_{i_s} + 1)$;

Append $b_{i,i+1}$ to $B$;

**end for**

**return** $C$ and $B$;

---

**Function** BIASRATEVECTOR $(G, L, L_g, L_{iter})$ **begin**

**Initialize** $u$ **and** $v$ as empty arrays;

$strand$ ← the strand of $G$;

$n \leftarrow$ size of $G$;

$n' \leftarrow n$;

$tag \leftarrow 0$;

$m \leftarrow |G|$;

**for** $i \leftarrow 1$ **to** $n$ **do**

    **if** $i > 1$ **then**

        $G \leftarrow G - \{g_1\}$;

        $G \leftarrow G - \{g_{1,2}\}$;

        $n' \leftarrow n' - 1$;

        $tag \leftarrow tag + 1$;

        $m = |G|$;

        $L \leftarrow$ update $L$ for $G$;

        $L_g \leftarrow$ update $L_g$ for $G$;

        $L_{inter} \leftarrow$ update $L_{inter}$ for $G$;

    **end if**

    **if** $strand = forward$ **then**

        **for** $j \leftarrow 1$ **to** $m$ **do**

            $x_j = 10^3 \times (l_m - l_{m-i+1}) \div (max_l - l_1)$;

        **end for**

        **for** $k \leftarrow 1$ **to** $n'$ **do**

            $sum \leftarrow 0$;

            **for** $t \leftarrow m - k_q + 1$ **to** $m - k_p + 1$ **do**

$sum = sum + f(x_t)$;

**end for**

$j = tag + k$;

$u_{i,j} = sum/\left(x_{m-k_p+1} - x_{m-k_q+1}\right)$;

Append $u_{i,j}$ to $u$;

**end for**

**if** $n' > 1$ **then**

**for** $k \leftarrow 1$ **to** $n' - 1$ **do**

$sum \leftarrow 0$;

**for** $t \leftarrow m - k_e + 1$ **to** $m - k_s + 1$ **do**

$sum = sum + f(x_t)$;

**end for**

$j = tag + k + 1$;

$v_{i,j} = sum/(x_{m-k_s+1} - x_{m-k_e+1})$;

Append $v_{i,j}$ to $v$;

**end for**

**end if**

**end if**

**if** $strand = reverse$ **then**

**for** $j \leftarrow 1$ **to** $m$ **do**

$x_j = 10^3 \times (l_i - l_1) \div (max_l - l_1)$;

**end for**

**for** $k \leftarrow 1$ **to** $n'$ **do**

$sum \leftarrow 0$;

**for** $t \leftarrow k_p$ **to** $k_q$ **do**

$sum = sum + f(x_t)$;

**end for**

$j = tag + k$;

$u_{i,j} = sum/\left(x_{k_q} - x_{k_p} + 1\right)$;

Append $u_{i,j}$ to $u$;

**end for**

**if** $n' > 1$ **then**

**for** $k \leftarrow 1$ **to** $n' - 1$ **do**

$sum \leftarrow 0$;

**for** $t \leftarrow k_s$ **to** $k_e$ **do**

$sum = sum + f(x_t)$;

**end for**

$j = tag + k + 1$;

$v_{i,j} = sum/(x_{k_e} - x_{k_s} + 1)$;

Append $v_{i,j}$ to $v$

**end for**

**end if**

**end if**

**end for**

**return** $u, v$;

**Table S2. Results of R-square statistic.** $R^2$ of the four bias rate functions $f_1(x) = 0.251319e^{0.001298x}$, $f_2(x) = 0.262360e^{0.001245x}$, $f_3(x) = 0.255069e^{0.001291x}$, and $f_4(x) = 0.253783e^{0.001286x}$.

| $R^2$ | $f_1(x)$ | $f_2(x)$ | $f_3(x)$ | $f_4(x)$ |
|---|---|---|---|---|
| $f_1(x)$ | 1 | 0.9981069 | 0.9991544 | 0.9999084 |
| $f_2(x)$ | 0.9982155 | 1 | 0.9987902 | 0.9989124 |
| $f_3(x)$ | 0.9991454 | 0.9987027 | 1 | 0.9993675 |
| $f_4(x)$ | 0.9999096 | 0.9988611 | 0.9993823 | 1 |

**Table S3. Results of predicted ATUs verified by experimental TSSs or promoters.** Overview of the four experimental TSS/promoter datasets (dataset 5 to 8) and the proportion of 5'-end genes and no 5'-end genes of the predicted ATUs by SeqATU for M9Enrich_Seq and RiEnrich_Seq, which were validated by experimental TSSs or promoters.

| | | dataset 5 | dataset 6 | dataset 7 | dataset 8 |
|---|---|---|---|---|---|
| **Source** | | Etwiller *et al.* (*13*) | RegulonDB TSSs | Thomason *et al.* (*14*) | RegulonDB promoters |
| **Technique** | | Cappable-seq | Collection | dRNA-seq | Collection |
| **TSSs/Promoters** | | 16,359 | 7,002 | 14,860 | 8,623 |
| **M9Enrich_Seq** | **5'-end genes** | 82% | 55% | 86% | 83% |
| | **no 5'-end genes** | 61% | 40% | 68% | 61% |
| **RiEnrich_Seq** | **5'-end genes** | 83% | 58% | 87% | 86% |
| | **no 5'-end genes** | 63% | 42% | 68% | 62% |

**Table S4. Results of predicted ATUs verified by computationally predicted TTSs.** Overview of the two computationally predicted TTS datasets (dataset 8 to 9) and the proportion of 3'-end genes and no 3'-end genes of the predicted ATUs by SeqATU for M9Enrich_Seq and RiEnrich_Seq, which were validated by experimental TTSs.

| | dataset 9 | dataset 10 |
|---|---|---|
| **Source** | Nadiras *et al.* (*40*) | Kingsford *et al.* (*41*) |
| **Type of TTSs** | Rho-dependent | Rho-independent |

| | | | |
|---|---|---|---|
| **TTSs** | | 7,806 | 2,630 |
| **M9Enrich_Seq** | **3’-end genes** | 79% | 63% |
| | **no 3’-end genes** | 59% | 29% |
| **RiEnrich_Seq** | **3’-end genes** | 79% | 62% |
| | **no 3’-end genes** | 58% | 29% |
